# Inter-individual and inter-regional variability of breast milk antibody reactivity to bacterial lipopolysaccharides

**DOI:** 10.1101/2024.05.16.594523

**Authors:** Lisa Crone, Jens Sobek, Nicole Müller, Tanja Restin, Dirk Bassler, Daniela Paganini, Michael B. Zimmermann, Patricia Zarnovican, Françoise H. Routier, Tais Romero-Uruñuela, Luis Izquierdo, Thierry Hennet

## Abstract

Breast milk is a vital source of nutrients, prebiotics, and protective factors, including antibodies and antimicrobial proteins. Using bacterial lipopolysaccharide arrays, we investigated the reactivity and specificity of breast milk antibodies towards microbial antigens, comparing samples from rural Kenya and urban Switzerland. Results showed considerable variability in antibody reactivity both within and between these locations. Kenyan breast milk demonstrated broad reactivity to bacterial lipopolysaccharides, likely due to increased microbial exposure. Antibodies primarily recognized the O-antigens of lipopolysaccharides and showed strong binding to specific carbohydrate motifs. Notably, antibodies against specific *Escherichia coli* O-antigens showed cross-reactivity with parasitic pathogens like *Leishmania major* and *Plasmodium falciparum*, thus showing that antibodies reacting against lipopolysaccharide O-antigens can recognize a wide range of antigens beyond bacteria. The observed diversity in antigen recognition highlights the significance of breast milk in safeguarding infants from infections, particularly those prevalent in specific geographic regions. The findings also offer insights for potential immunobiotic strategies to augment natural antibody-mediated defense against diverse pathogens.

## INTRODUCTION

Breast milk supplies infants with crucial nutrients, prebiotic elements such as complex oligosaccharides, and protective components such as antibodies and antimicrobial proteins (1). The composition of breast milk evolves throughout lactation. Colostrum, which is only secreted within the first days after birth, contains the highest concentration of antibodies (2, 3). These antibodies, which predominantly belong to the IgA isotype, play a critical role in protecting the infant’s gastrointestinal tract against infections (4). Breast milk antibodies originate from maternal B-cells that mature in the gastrointestinal tract and migrate to the lactating mammary gland (5). Exposure to bacterial antigens in the gut stimulates these B-cells, leading to the production of antibodies secreted into breast milk. This process ensures that the milk contains diverse antibodies tailored to the mother’s gut microbiota, providing the infant with customized protection against potential pathogens (4, 6, 7). Breast milk antibodies work alongside maternal IgG antibodies transferred to the fetus during pregnancy, offering a comprehensive defense against potential infections (8). IgA antibodies in breast milk are particularly effective at neutralizing pathogens in the infant’s gastrointestinal tract, a common site of infections in newborns. Additionally, breast milk antibodies can recognize and bind to numerous pathogens, preventing their attachment to host cells and subsequent invasion. Overall, breast milk antibodies are critical components of an infant’s immune system and play an essential role in promoting their health and development (9–11).

Maternal B-cells in the mammary gland are primed to produce antibodies that recognize a wide range of antigens presented by gut microbes (12). These microbes, mainly gram-positive and -negative bacteria, are covered with carbohydrate antigens like peptidoglycan, teichoic acid, and lipopolysaccharide (LPS). Especially the large diversity of LPS O-antigens on gut bacteria represents a vast repertoire of carbohydrate antigens that stimulate the immune system (13). The precise mechanisms by which carbohydrate antigens stimulate B-cells and mediate Ig class switching in the gut are still being studied. In mucosal tissues, such as the intestinal lamina propria, B-cell activation and immunoglobulin class switching can occur independently of T-cells through the stimulatory effects of the TNF superfamily proteins BAFF and APRIL secreted by dendritic cells (14, 15). Because of the constant exposure of mucosal immune cells to gut bacteria, bacterial carbohydrate antigens are major stimuli for antibody production, thus leading to high titers of carbohydrate-specific antibodies in the gastrointestinal tract and blood circulation. Carbohydrate structures, which are prominently exposed on bacterial surfaces but absent on animal cells, elicit robust antibody responses. These carbohydrate antigens mainly consist of monosaccharides, such as rhamnose and galactofuranose (Gal*f*), and disaccharides, such as the Gal(α1-3)Gal α-Gal antigen and GalNAc(α1-3)GalNAc Forssman antigen, which are not found on human cells. Antibodies directed towards ABO blood group antigens are also generated through the stimulation of B-cells by structurally similar carbohydrates exposed on gut bacteria (16).

Despite significant differences between prokaryotic and eukaryotic glycosylation pathways, specific carbohydrate structures are shared across taxonomic groups. Antibodies targeting bacterial glycans may therefore cross-react with similar epitopes presented on other organisms. For example, the presence of *E. coli* O86 in the gut stimulates the production of antibodies reacting with the a-Gal epitope occurring on its O-antigen, and which is also found on the surface of pathogens such as *Plasmodium falciparum*, thereby conferring protection towards malaria (17). However, the production of cross-reactive antibodies may also promote autoimmune responses when the recognized carbohydrate epitopes are also presented on host cells. Infection with *Campylobacter jejuni* leads to antibodies recognizing sialylated lipooligosaccharides on the bacterial surface, which resemble GM1 gangliosides presented on peripheral nerves. The production of antibodies cross-reacting with host gangliosides are associated with the development of Guillain-Barré syndrome characterized by immune-mediated peripheral neuropathy (18).

Although our understanding of the significance of breast milk antibodies has progressed, many questions remain unanswered. The range of antigens recognized by breast milk antibodies is not yet fully understood, nor is the functional relevance of cross-reactive protection. The specific repertoire of carbohydrate antigens recognized by milk antibodies also remains unknown, and it is unclear whether these antigens can cross-react with infectious pathogens relevant to different regions of the world. In our present study, we addressed these questions by examining the repertoire of antigens recognized by antibodies in breast milk collected from different geographical locations. Using an array of LPS antigens, we aimed to better understand the specific composition of breast milk antibodies and how they vary between lactating mothers and across different populations. Our research highlights the critical role of carbohydrate-specific antibodies in providing protection against different pathogens, emphasizing the importance of cross-reactivity in conferring an effective immune protection.

## RESULTS

Multiple factors such as genetics, diet, gut microbiota composition and the environmental exposure to pathogens influence mucosal antibody production. Considering the role of these factors in shaping the repertoire of antibodies secreted in breast milk, we compared the reactivity of breast milk antibodies from two geographical locations towards a panel of bacterial LPS antigens. The O-antigens at the tip of LPS represents a vast diversity of carbohydrate antigenic structures, characterized by oligosaccharide repeats (19, 20). These oligosaccharides consist of di- to octasaccharides comprising monosaccharides found in eukaryotic glycans and a broad range of monosaccharides unique to prokaryotes (16, 21). For our LPS arrays, we have selected a panel of 103 LPS representing a broad range of O-antigens, some of them including carbohydrate epitopes found on protist and animal cells such as α-Gal and the Forssman antigens (Supplementary Table 1). Rough mutant LPS featuring truncated glycan chains were also included on the array to assess the specificity of breast milk antibody towards carbohydrate O-antigens.

Pure LPS (Supplementary Figure 1A) were first spotted at four concentrations on NEXTERION® 3-D Hydrogel coated slides to evaluate the correlation between the quantity of printed antigen and signal intensity relevant to antibody recognition of these antigens. We noted a direct proportionality in antibody binding with increasing LPS concentrations for both O-antigen-specific immunoglobulin G (IgG) and breast milk immunoglobulin A (IgA) (Supplementary Figure 1B and 1C), thus confirming the suitability of the method to estimate antibody specificity to the LPS recognized. The method also allows estimating binding strength when testing pure antibodies. When testing antibody mixtures, binding to LPS does not allow to discriminate between low total levels of antibodies with high affinity and high total levels of antibodies with generally low affinity towards LPS. Because of the mass heterogeneity of LPS related to the variable degree of O-antigen polymerization, LPS were printed in a weight to volume ratio and not at a specific molarity. Consequently, the signals obtained for antibody binding to the printed LPS are semi-quantitative.

To compare the antigenic specificity of breast milk antibodies originating from different geographical locations and environments, we investigated a cohort of breast milk samples collected from a rural region in Kenya (22) and breast milk samples collected from an urban environment in Switzerland. The average age of the donor mothers at delivery was 27 years for the Kenyan samples and 32 years for Swiss samples (Supplementary Table 2). In Kenyan breast milk, the concentrations of IgA, IgM, and IgG ranged from 0.25 to 0.74 mg/mL, 0.01 to 0.35 mg/mL and 0.02 to 0.10 mg/mL, respectively. In Swiss breast milk, the concentration range of IgA, IgM, and IgG were 0.24 to 1.47 mg/mL, 0.03 to 0.44 mg/mL and 0.01 to 0.15 mg/mL, respectively. IgM concentrations were higher in Swiss samples, whereas IgA and IgG concentrations were comparable in breast milk samples from both locations (Supplementary Figure 2). We first compared the reactivity of breast milk IgA, IgM and IgG from the 30 breast milk samples from Kenya towards the 103 LPS printed on the array. As expected, the strongest signals were measured for IgA, which is the most abundant antibody class in breast milk (Figure 1). Overall, a strong variability in the recognition of LPS antigens by IgA was observed within the breast milk samples tested. The strongest reactivities were detected towards LPS from the *E. coli* serotypes O13, O16, O55, O82, O85, O86, O111, O142, and K-235. The LPS of further Enterobacteriaceae including *Shigella flexneri* 2a and *Klebsiella pneumoniae* O1 were also strongly recognized, whereas low IgA reactivity was detected towards most of the *Citrobacter* LPS tested. The pattern of LPS recognized by IgA also significantly varied between individual breast milk samples, suggesting that LPS-specific IgA reflected previous exposure of the women’s immune system to bacteria presenting the corresponding epitopes. Given that IgM and IgG make up about 8% and 2% of antibodies in human breast milk, the reactivity of these Ig classes towards LPS was considerably weaker than the reactivity of IgA (Figure 1). When scanning the arrays at higher sensitivity, the signals measured for IgM and IgG binding pointed to differences in the recognition of individual LPS against IgA reactivity (Supplementary Figure 3), suggesting that individual LPS do not lead to the production of similar antibody responses across Ig classes. For all LPS investigated, the respective reactivity of IgA and IgM did not correlate significantly (Supplementary Figure 4). The same lack of correlation was observed between the reactivity of IgA and IgG towards individual LPS (Supplementary Figure 5).

**Figure 1.**
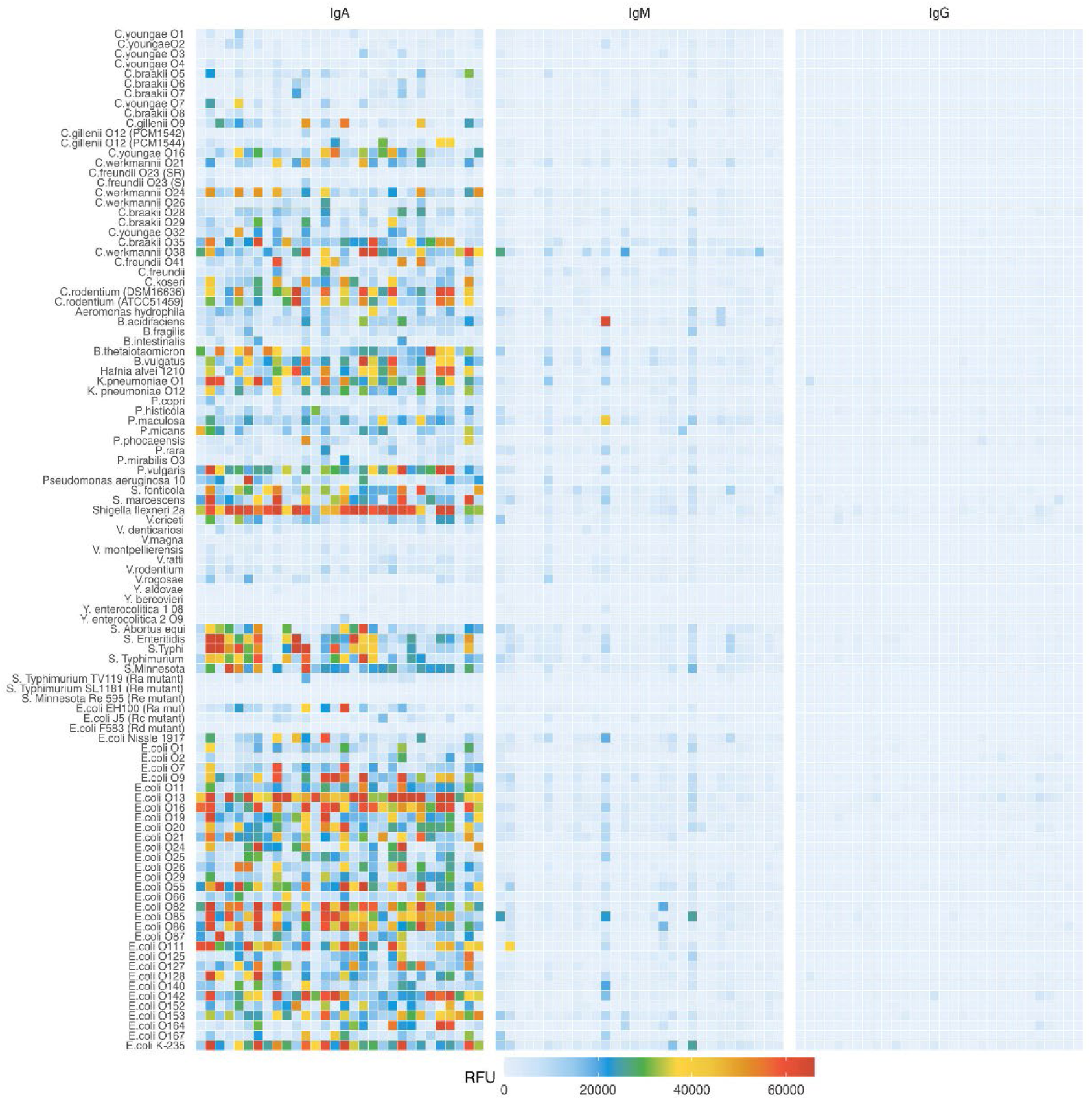
IgA, IgM and IgG reactivity to LPS in Kenyan breast milk samples. Breast milk samples were normalized to an IgA concentration of 10 μg/ml and applied to LPS arrays. Antibody binding to LPS was detected using either an anti-human IgA and IgM Cy3 or anti-human IgG AlexaFluor647 labelled antibody. Fluorescence signals were recorded at a PMT-gain of 190V. Data are shown as net mean RFU values of 9-replicate dots.

The incorporation of rough mutant LPS on the array allowed assessing which LPS moiety was mainly recognized by antibodies. A series of mutant LPS lacking only the O-antigen (Ra) or the O-antigen plus portions of the core oligosaccharide (Rc, Rd, Re, Figure 2A) confirmed that the O-antigen was the main epitope recognized by breast milk IgA, IgM and IgG. Reactivity towards the Ra and Rc mutant LPS of *E. coli* tested was on average 10-fold lower than towards smooth LPS featuring O-antigens (Figure 2B and 2C). Similar differences were measured between the smooth LPS from *Salmonella enterica* containing O-antigens and the rough mutant LPS lacking portions of the carbohydrate moiety (Figure 2D and 2E).

**Figure 2.**
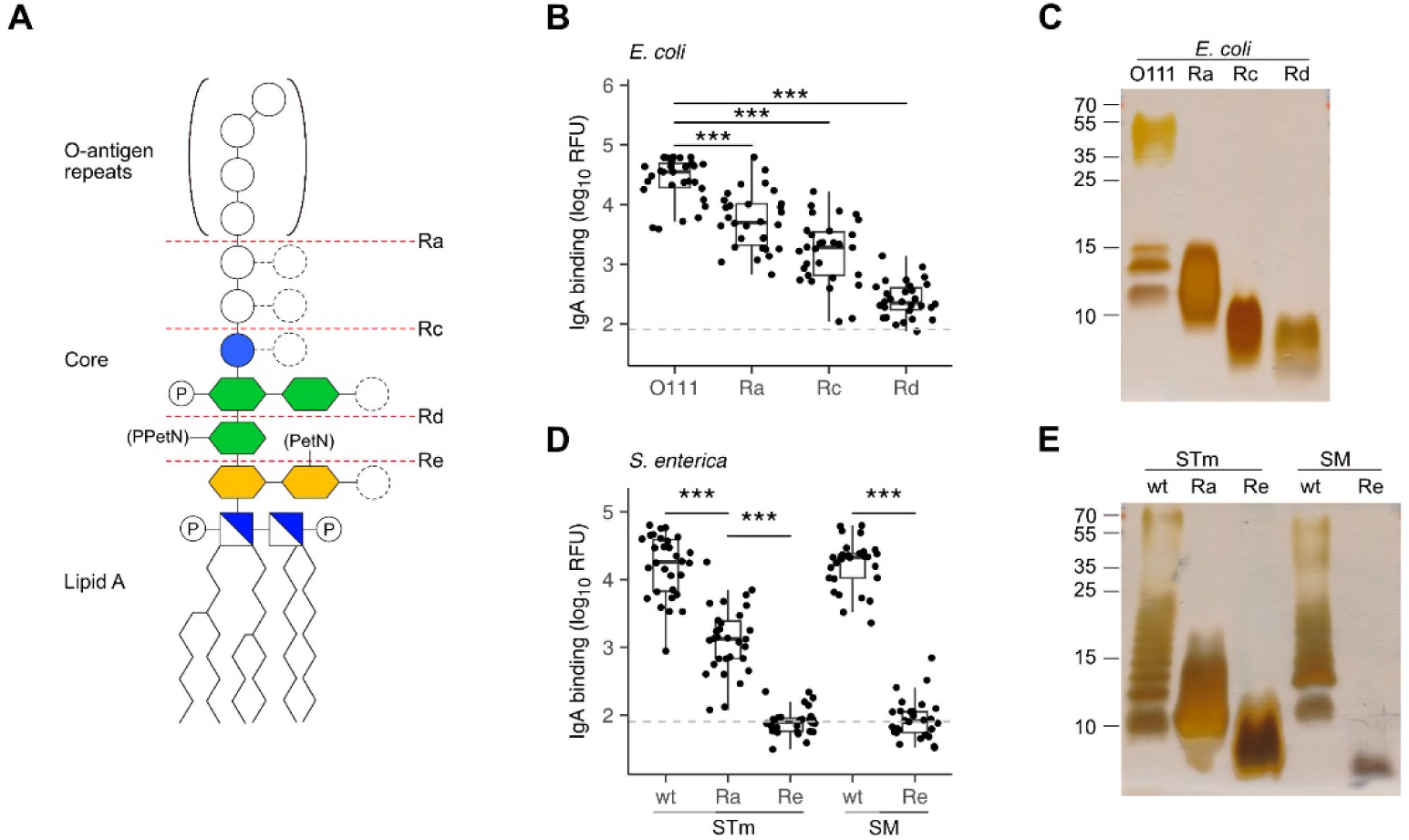
Recognition of O-antigens by breast milk IgA. (A) Schematic structure of LPS consisting of lipid A, a phosphorylated (P) glucosamine disaccharide (blue/white square), a glycan core of 2-keto-3-deoxy-D-mannooctanoic acid (KDO, yellow hexagon), heptose (green hexagon), glucose (blue circle) and repeats of O-antigen oligosaccharides. Distinct serotypes vary in sugar composition (empty circles) and number of sugars (dashed circles) and can be modified with phosphoethanolamine (PetN) or pyrophosphorylethanolamine (PPEtN). *E. coli* and *S. enterica* rough mutants lack the O-antigen (Ra) or variable regions of the glycan core (Rc, Rd and Re). (B) IgA binding intensity to *E. coli* smooth LPS (O111) and rough LPS (Ra, Rc, Rd). (C) Silver stain of *E. coli* smooth LPS (O111) and rough LPS mutants (Ra, Rc, Rd) separated by SDS-PAGE. (D) IgA binding intensity to *S. enterica* Typhimurium (STm) and S. Minnesota (SM) wildtype (wt) smooth LPS and Ra, Re rough LPS mutants. (E) Silver stain of STm and SM wt smooth LPS and rough LPS (Ra, Re) separated by SDS-PAGE. P values were determined by unpaired Wilcoxon test and indicated by asterisks (***, p < 0.001). The grey dotted line marks the threshold of detection (TOD).

### Comparison of antibody reactivity between Switzerland and Kenya

The analysis of breast milk samples from Kenya revealed a wide inter-individual variability in the reactivity of breast milk antibodies to LPS (Figure 1). To evaluate the geographical specificity of the breast milk antibody response to LPS antigens, we have also analyzed the antibody reactivity of breast milk samples obtained from an urban environment in Switzerland. Generally, stronger and broader IgA reactivities to LPS were detected in the breast milk samples from Kenya (Figure 3). In particular, we detected higher IgA reactivities in Kenyan samples towards the LPS of bacteria causing diarrheal disease, such as *E. coli*, *S. flexneri*, and *Salmonella enterica*. We also observed that LPS containing specific mono- and disaccharide motifs, such as rhamnose, 3,6-dideoxysugars, Gal*f*, α-Gal and Forssman epitopes resulted in strong IgA binding in Kenyan breast milk samples (Figure 4, Supplementary Figure 6). Rhamnose is a carbohydrate widely occurring in plants (23) and bacterial cell walls (24) but absent in animals (25), which explains the high titers of antibodies in humans recognizing rhamnose. The O-antigens of *S. flexneri* 2a and *E. coli* O13 (Figure 4A and 4B) include three a-linked rhamnose residues. Interestingly, the O-antigen of *Citrobacter gillenii* O9, which is made up of a-linked N-acetyl-rhamnosamine (RhaNAc), was about ten-fold less recognized than the rhamnosylated O-antigen of *S. flexneri* 2a and *E. coli* O13 (Figure 4C). Recognition of *E. coli* O86, which contains the a-Gal and Forssman disaccharide motifs was stronger for IgA from Kenyan samples than Swiss samples (Figure 4D). IgA reactivity towards LPS containing the 3,6-dideoxy sugars abequose, tyvelose and colitose was also stronger in Kenyan samples than Swiss samples (Figure 4E and 4F, Supplementary Figure 6D-G). These three 3,6-dideoxy sugars are commonly found in *Salmonella enterica* subsp. *enterica* serovar *S.* Typhimurium and *S.* Typhi, which are endemic in developing countries, cause non-typhoidal salmonellosis, and typhoidal fever, respectively (26). *Salmonella enterica* serovar Typhimurium (Figure 4E) and serovar Abortus equi (D) share similar O-antigens including abequose, while *Salmonella enterica* serovar Typhi shares a similar O-antigen but with tyvelose instead of abequose (Figure 4F). Abequose is also found in the O-antigen of *C. werkmannii* O38, which was strongly recognized by IgA in Kenyan samples (Supplementary Figure 6E). Colitose occurs in *E. coli* O55 and *E. coli* O111, which both elicited strong IgA responses in Kenyan samples (Supplementary Figure 6F and G). The enteropathogenic *E. coli* O111 serogroup is the primary cause of diarrhea in infants, which is particularly a problem in developing countries (27). Further LPS that were predominantly recognized by IgA in Kenyan breast milk samples were *E. coli* O86, O87, O127 and O142, which contain the a-Gal and/or Forssman disaccharide antigens (Figure 4D, Supplementary Figure 6H, I and L). Humans produce high antibody titres towards these carbohydrate antigens, which are absent on human cells (28, 29). Also, LPS containing Gal*f*, a monosaccharide that is not found in humans (30) were strongly recognized by IgA in Kenyan samples (Supplementary Figure 6J-L). Altogether, Kenyan breast milk samples featured higher IgA reactivities towards LPS associated with disease in developing countries. The presence of such reactive antibodies in the breast milk certainly contributes to the protection of nursed infants towards a broad range of bacterial pathogens.

**Figure 3.**
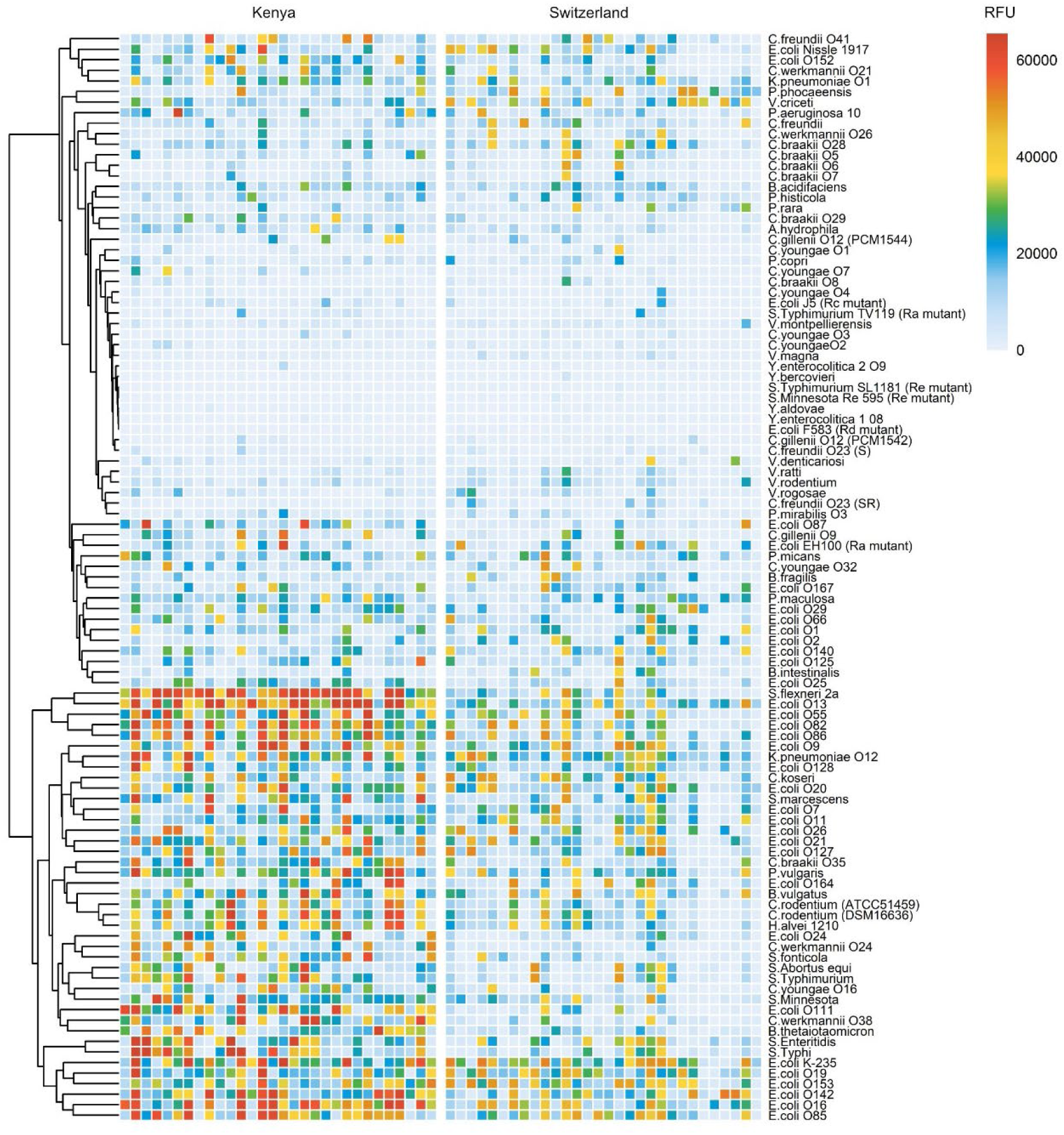
Comparison of IgA reactivity between Kenyan and Swiss breast milk samples. 30 breast milk samples from Kenya (left panel) and 30 samples from Switzerland (right panel) were normalized to an IgA concentration of 10 μ g/ml and analyzed on LPS arrays. Antibody binding to LPS was detected using anti-human IgA secondary antibody coupled to Cy3. Fluorescence signals were recorded at a PMT-gain of 190V. The net mean RFU values of 9-replicate dots are shown. Hierarchical clustering of LPS binding signals was performed using the pheatmap package in R studio.

**Figure 4.**
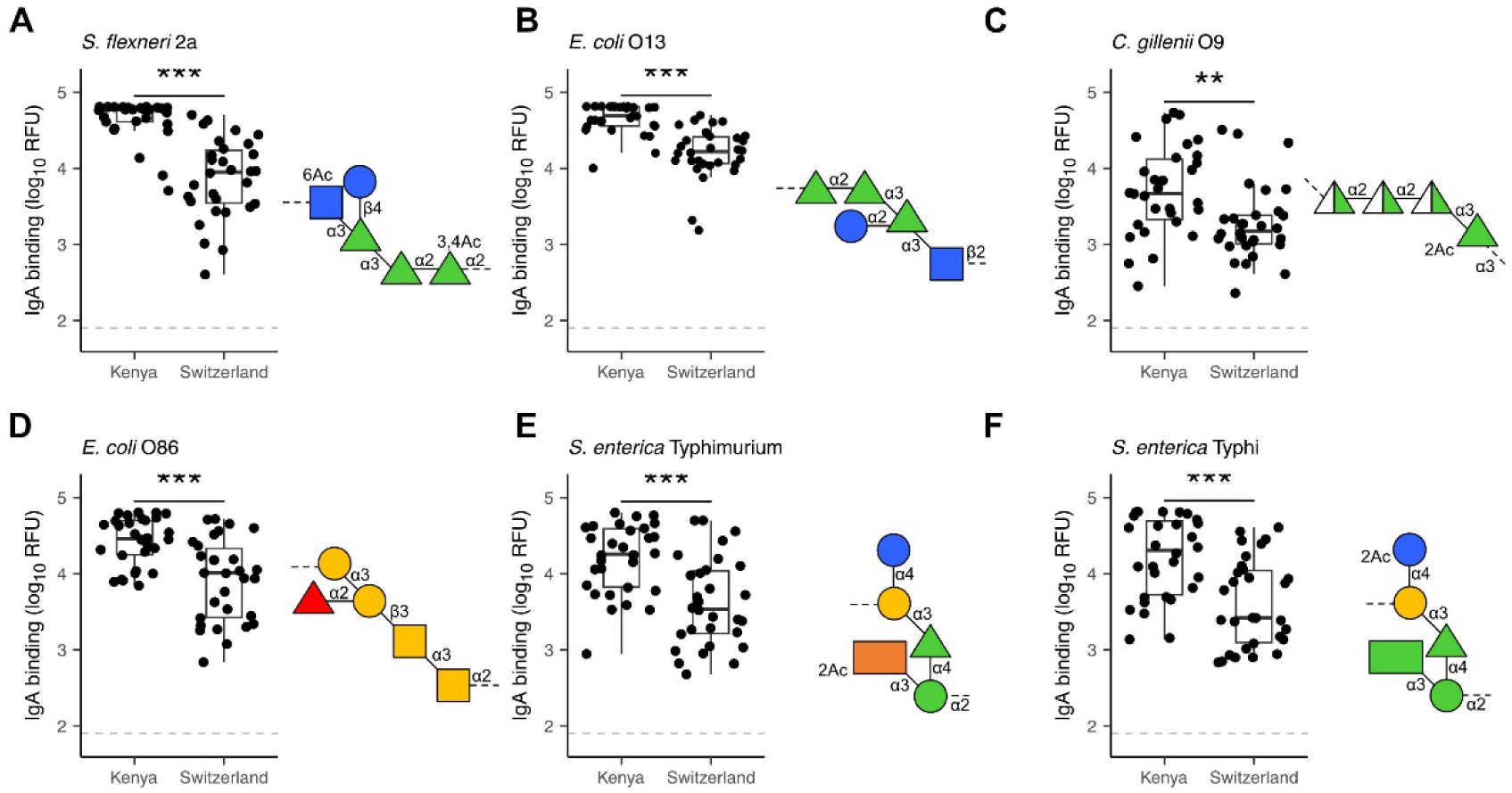
Differential recognition of specific LPS O-antigens by Kenyan and Swiss breast milk IgA. IgA binding to LPS O-antigens including the immunogenic epitopes rhamnose (green triangles) as found in (A) *S. flexneri* 2a; (B) *E. coli* O13 and (C) *C. gillenii* O9; the α-Gal (two yellow circles) and Forssman (two yellow squares) antigens as found in (D) *E. coli* O86; the dideoxy sugars abequose (orange rectangle) and tyvelose (green rectangle) as found in (E) *S. enterica* Typhimurium and (F) *S. enterica* Typhi. IgA binding was detected using anti-human IgA Cy3, scanned at a PMT-gain of 190V. The grey dotted line shows TOD. Comparisons of net mean RFU values between Kenyan (n=30) and Swiss (n=30) samples was done using an unpaired Wilcoxon test. P-values <0.001 (***), <0.01 (**) were considered statistically significant. The symbols used to display O-antigens is based on the standard symbolic nomenclature for glycans (50).

### Cross-reactivity of LPS-specific antibodies

The vast diversity of O-antigens not only encompasses a multitude of distinct antigens but often gives rise to structurally similar antigens. Hierarchical clustering of our array analysis pointed to similar antibody responses towards LPS sharing similar O-antigen structures. For example, the similar O-antigens of *E. coli* O13 and *S. flexneri* 2a (Figure 4A and 4B) elicited cross-reactive antibody responses among the Kenyan breast milk samples analyzed (Figure 3). Likewise, the structurally overlapping O-antigens of *S*. Enteritis and *S*. Typhi (Figure 4F) were similarly recognized across the breast milk samples analyzed (Figure 3), thus suggesting recognition by cross-reacting antibodies. To confirm the cross-reactivity of antibodies between structurally similar O-antigens, we have analyzed the reactivity of a polyclonal antibody specific to the *E. coli* O-antigen O152. The same O-antigen is also expressed by *Citrobacter rodentium* ATCC51459 and similar O-antigens are found on *Hafnia alvei* 1210 and *Klebsiella pneumoniae* O12, which both contain the rhamnose(b1-4)N-acetylglucosamine (GlcNAc) motif (Figure 5A). Using immunoblotting, we confirmed that the anti-O152 antibody indeed recognized the purified LPS from *C. rodentium* and *H. alvei* (Figure 5B). The LPS of *K. pneumoniae* O12 remained undetected by the anti-O152 antibody, which may be due to the masking of b-rhamnose by the direct linkage to the next GlcNAc unit, which may prevent the recognition by the antibody.

**Figure 5.**
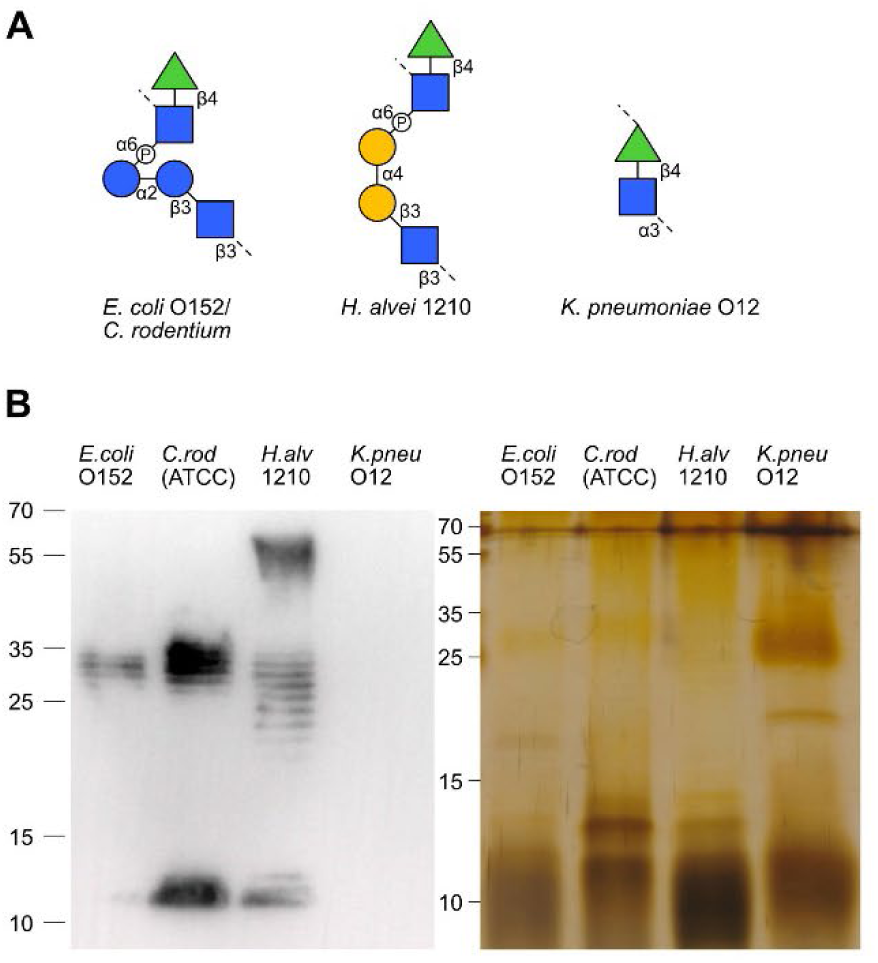
Antibody cross-reactivity to structurally similar LPS O-antigens. (A) The O-antigen structures of *E. coli* O152/*C. rodentium* (ATCC51459), *H. alvei* 1210 and *K. pneumoniae* O12. (B) Immunoblotting and silver stained SDS-PAGE of *E. coli* O152, *C. rodentium*, *H.alvei* 1210 and *K. pneumoniae* O12 LPS recognized by anti-O152 polyclonal antibody.

Beyond cross-reactivity between O-antigens, antibodies recognizing bacterial carbohydrate antigens likely also recognize similar carbohydrate epitopes presented on eukaryotic cells. Previous studies have demonstrated the ability of antibodies targeting bacterial antigens to cross-react with parasitic antigens. The detection of strong IgA responses towards α-Gal and Gal*f*-containing O-antigens in our analysis suggested that breast milk antibodies against these epitopes may cross-react with similar epitopes on parasitic pathogens endemic in Kenya, such as *Plasmodium falciparum* (31) and *Leishmania major* (32). To test this hypothesis, we have investigated the reactivity of O-antigen-specific antibodies to protein extracts of *P. falciparum* and *L. major* using immunofluorescence and immunoblotting. Antibodies specific to *E. coli* O142 strongly bound to blood stage *P. falciparum*, as well as weaker bindings by *E. coli* O152 (Figure 6) which is consistent with the results from immunoblotting, where they weakly reacted with an antigen of *P. falciparum* at asexual intraerythrocytic stages (Figure 6E and 6F). *E. coli* O55- and O85-specific antibodies poorly bound to *P. falciparum* (Figure 6). Antibodies specific to *E. coli* O55, O85, O142 and O152 were found to bind to *L. major* at the promastigote stage to various extent as detected by immunofluorescence (Figure 6B). We also tested reactivity of these O-antigen-specific antibodies to *L. major* wildtype and to a mutant strain lacking Gal*f* (33) by Western blotting. The anti-O85 antibody recognized an antigen on both wildtype and Gal*f* -deficient mutant *L. major* (Figure 6D), which was unexpected considering the presence of Gal*f* in the O85 O-antigen. The anti-O142 antibody did not reveal any antigen binding of *L. major* on Western blot, whereas the *E. coli* O152 antibody recognized several antigens of *L. major* extracts (Figure 6D). By comparison, the anti-O55 antibody failed to cross-react with any antigen of *P. falciparum* and *L. major* on Western blot (Figure 6C) despite its strong binding to *L. major* detected by immunofluorescence (Figure 6B). The confirmation that O-antigen-specific antibodies indeed recognized epitopes on parasite extracts indicate that the vast repertoire of carbohydrate-specific cross-reactive antibodies in the breast milk could provide a broad protection of nursed infants, not just towards bacterial infections but also towards other endemic eukaryotic pathogens.

**Figure 6.**
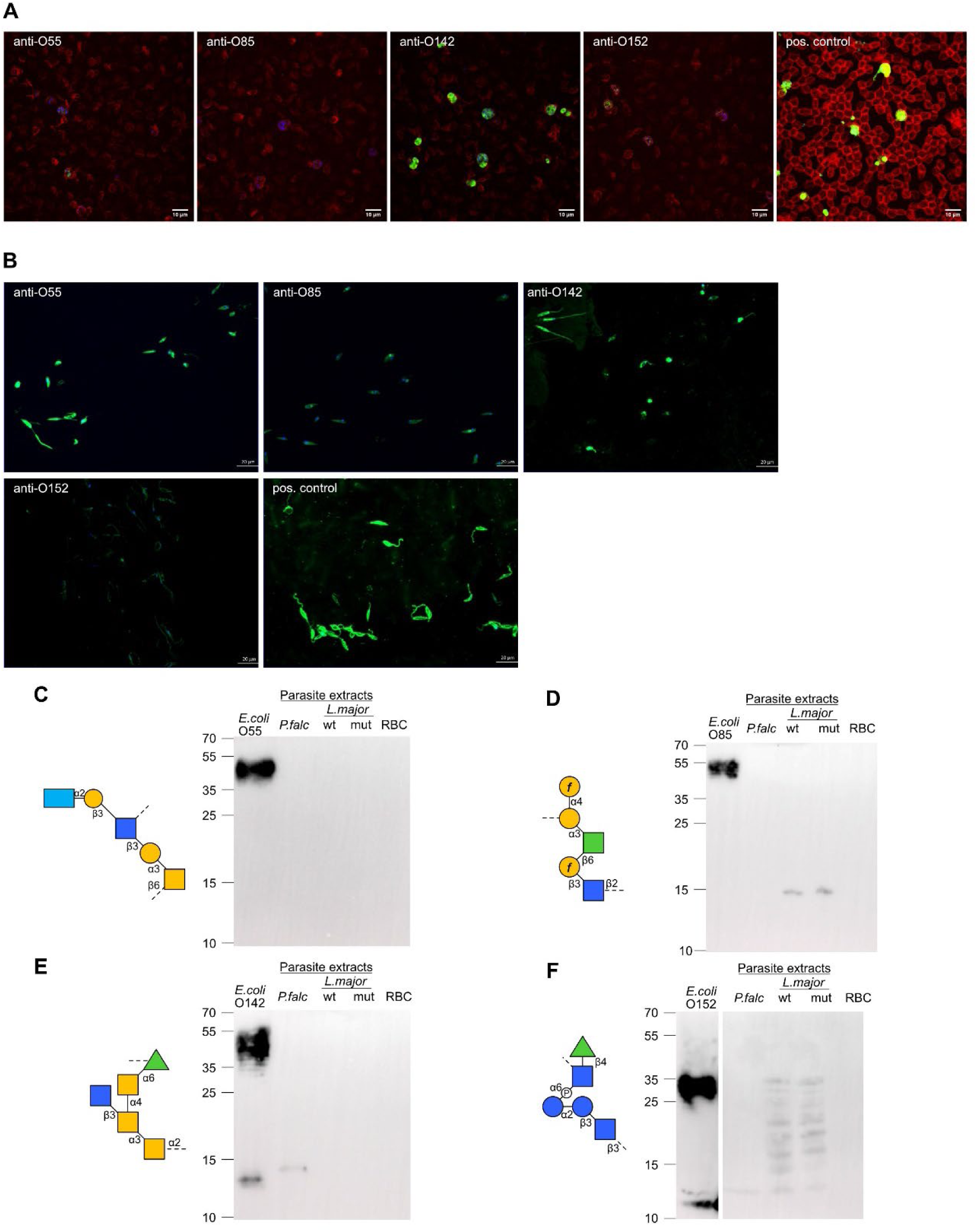
Cross-reactivity of O-antigen-specific antibodies to *L. major* and *P. falciparum* antigens. (A) Immunofluorescence staining of *P. falciparum* non-synchronized cultures with anti-O-antigen antibodies specific towards *E. coli* O55, O85, O142, O152 and with positive control antibody. (B) Immunofluorescence staining of *L. major* promastigotes with anti-O-antigen antibodies specific towards *E. coli* O55, O85, O142, O152 and with positive control antibody. (C-F) SDS-PAGE with *E. coli* LPS, parasite extracts from *P. falciparum* (*P.falc*) and *L. major* wildtype (wt) and Gal*f* mutant (mut), and red blood cell (RBC) control. Immunoblotting using O-antigen specific sera from (C) *E. coli* O85, (D) *E. coli* O142, (E) *E. coli* O152 and (F) *E. coli* O55.

## DISCUSSION

Our investigation of breast milk antibodies not only underlined the broad range of bacterial LPS antigens recognized, but also the evident variability in the reactivity of these antibodies between individual breast milk samples. The inter-individual variability in the reactivity of breast milk IgA toward bacterial antigens has been reported in previous studies (34, 35), which also pointed to differences in the recognition of bacteria by breast milk antibodies collected from various geographical locations (34). These studies however only surveyed few bacteria without discrimination of the antigens recognized. The application of an array of pure LPS antigens provided here a more precise mapping of the range and types of bacterial epitopes recognized by breast milk IgA, IgM and IgG. The resulting higher resolution of antibody specificities confirmed the variability of antigens recognized between individual breast milk samples and between samples collected in different geographical locations. These differences in antigen recognition likely reflect the microbial exposure of the mothers prior to lactation, as interactions of microbes with mucosal immunity in the gut are known to shape the range and amount of IgA secreted in the breast milk (36, 37).

The broader range of LPS antigens recognized and the stronger IgA binding measured for the breast milk samples from Kenya likely reflect the prominent microbial exposure of these mothers living in rural regions with limited access to clean water, which increases their susceptibility to gastrointestinal infections. Whereas the elevated infection pressure represents a significant health risk especially for women during pregnancy, the increased exposure to microbes obviously leads to the generation of breast milk that is rich in protective antibodies towards a vast range of bacterial pathogens. The broad spectrum of breast milk antibodies against microbes therefore underscores the importance of breastfeeding to ensure the protection of infants, especially in locations with precarious environmental health conditions.

The application of pure LPS structures on the antigen arrays also allowed defining the epitopes that are mainly recognized by breast milk antibodies. The presence of rough mutant LPS on the array, which lack portions of the carbohydrate moiety, clearly showed that breast milk IgA, IgM and IgG predominantly recognize O-antigens. Re rough mutants featuring lipid A with only two KDO residues were hardly recognized by breast milk antibodies. Studies on the specificity of monoclonal antibodies cloned from intestinal IgA and lactating mammary gland IgA pointed to the polyreactivity of these antibodies to various microbial antigens such as LPS, capsular polysaccharides and flagellin (38). The polyreactivity of these IgA monoclonal antibodies was shown through their ability to bind to multiple bacterial targets (12). Although the mechanisms of such a polyreactivity are not completely clear, the phenomenon can be explained by the sharing of similar glycan epitopes across multiple bacterial taxa. The relevance of glycan epitope similarity in eliciting polyreactive IgA and IgM was confirmed in the case of cross-specific protective antibodies targeting LPS of various serotypes of *Klebsiella pneumoniae* and even structurally-similar LPS of non–*Klebsiella pneumoniae* species (39). Beyond the example of *Klebsiella pneumoniae* LPS, our study shows that O-antigen similarity leads to the development of antibodies that confer a broad recognition of bacterial antigens, thereby providing protection against a wide range of invading pathogens. While our study demonstrates the ability of O-antigen specific antibodies to bind to *Plasmodium falciparum* and *Leishmania major* cells, the extent and functional significance of cross-reactivity of breast milk antibodies towards LPS and eukaryotic antigens still requires further investigation.

O-antigens including carbohydrate motifs that are absent from human cells elicited strong IgA binding, as especially evidenced in breast milk samples from Kenya. Monosaccharides, such as rhamnose, colitose, abequose, tyvelose, Gal*f* and the disaccharides α-Gal and Forssman antigens were associated with the strongest signals recorded for IgA binding on the LPS array. Some of these immunogenic carbohydrate antigens do not only occur on bacterial polysaccharides but are also found on eukaryotic cells including protists, fungi plants and animals. The 3,6-dideoxysugar tyvelose is part of an immunodominant surface antigen of the nematode *Trichinella spiralis* (40). Gal*f* is also commonly found on of the glycans of the kinetoplastids *Leishmania spp.* (41) and *Trypanosoma spp.* (42). The α-Gal antigen also occurs on *P. falciparum* (17), while the presence of antibodies to this carbohydrate antigen has been correlated with protection against malaria transmission (43, 44). Interestingly, gut colonization with *E. coli* O86, which contains α-Gal antigen in its O-antigen, was shown to be associated with malaria protection in humans and in a mouse model (17). This protection was related to the production of anti-α-Gal IgM, whereas IgG to α-Gal did not appear to confer protection against *P. falciparum* infection. Beyond *E. coli* O86 colonization and antibodies towards the α-Gal antigen, our present work shows that further *E. coli* serotypes stimulate the production of antibodies, which cross-react with *P. falciparum* and *L. major* antigens. This cross-reactivity of anti-LPS antibodies to parasites indicates that the *E. coli* O-antigens may stimulate the production of antibodies conferring partial protection to malaria and leishmaniasis. In general, commensal bacteria featuring carbohydrate antigens shared with specific pathogens could be used in immunobiotic approaches to boost the natural antibody protection towards a broad range of pathogens, which would be especially relevant for infections where vaccines are still unavailable.

In conclusion, breast milk antibodies targeting carbohydrate antigens on bacteria may have a protective effect against a wide range of pathogens beyond prokaryotes. In the context of Kenya, where endemic infections such as malaria and leishmaniasis impose significant burden on local populations, carbohydrate-reactive breast milk antibodies may contribute to protect infants from multiple infections.

## METHODS

### Breast milk samples

Breast milk samples were collected at the University Hospital Zurich, Switzerland. Breast milk samples from Kenya were collected in a hospital and health center in Kwale County, southern coastal Kenya (22) (Supplementary Table 2). Ethical approval was given by the Ethical review committees of ETH Zurich (EK 2019-N-59) and Jomo Kenyatta University of Agriculture and Technology, Nairobi, Kenya (JKU/2/4/896B). IgA, IgM and IgG levels in breast milk samples were measured using commercially available ELISA kits (Abnova, Taiwan) according to the manufacturer’s recommended protocols.

### Bacterial cultures

Pure bacterial strains were purchased from the Leibniz Institute German Collection of Microorganisms and Cell Cultures GmbH (DSMZ, Braunschweig, Germany), the Polish Collection of Microorganisms (PCM, Wroclaw, Poland), the National Collection of Type Cultures (NCTC, Salisbury, UK) and the American Type Culture Collection (ATCC, Manassas, USA). Bacteria were cultured under conditions recommended by the respective repositories (Supplementary Table 1). Obligate anaerobic bacteria were grown in an anaerobic workstation (Bugbox, Baker, Sanford).

### LPS purification

LPS (S1 Table) was purified using the hot aqueous-phenol method (45). Briefly, bacteria were harvested from a 1 L culture at OD_600_ 2.0, pelleted by centrifugation at 8,000 x g for 10 min, washed with 40 mL of ice-cold water and pelleted again. Pellets were resuspended in 20 mL of 2 % SDS, 10 % glycerol, 2 % β-mercaptoethanol in 50 mM Tris-HCl, pH 6.8, boiled at 95 °C for 30 min, then incubated twice with a final concentration of 50 μg/mL DNase I (Roche, Basel, Switzerland) and RNase A (Roche, Basel, Switzerland) at 37 °C for 30 min, subsequently followed by another incubation with proteinase K (Thermo Scientific, Waltham, MA, USA) at a final concentration of 50 μg/mL at 55 °C overnight. Equal volumes of phenol (Sigma-Aldrich, St. Louis, USA) were added to the samples, which were incubated for 15 min at 65 °*C* after thorough mixing. A volume of 150 mL of diethyl ether was added, samples were vortexed for 30 s before centrifuging at 13,000 x g for 20 min. The upper phase was discarded, and phenol extraction was repeated using the lower phase. For precipitation, samples were supplemented with 0.5 M sodium acetate, along with 10-times the volume of 95 % ethanol and were kept overnight at −20 °*C.* After centrifugation at 3,000 x g at 4 °C for 10 min, the supernatant was discarded. The pellet was resuspended in 20 mL deionized water and dialyzed using the Pur-A-Lyzer Mega 3500 Dialysis Kit (Sigma-Aldrich) and lyophilized. To ensure purity of the LPS fractions, phenol extractions and dialysis were repeated. Final products were stored at −20 °C.

### LPS arrays

LPS stocks were diluted to 250, 125, 62 and 31 μg/mL in 0.5 M phosphate buffer at pH 8.and spotted to NEXTERION® 3-D Hydrogel coated slides (Schott Nexterion, Jena, Germany) using a sciFLEXARRAYER S11 microarray spotter (Scienion, Berlin, Germany). For immobilization, the slides were stored for one to six days in a humidity chamber containing a saturated sodium chloride solution, setting 75 % relative humidity. All following processing steps were performed in an automated hybridization station (HS4800, Tecan, Männedorf, Switzerland), which allows to control temperature and duration of all incubation and washing steps. The incubations with 40 mg/mL Blotting-Grade Blocker (BioRad, Hercules, CA, USA) as blocking buffer, breast milk samples, and secondary antibody were followed by washing steps consisting of a rinse for 1 min with PBS-T followed by an agitation of the stationary solution for 3 min. Typically, series of up to 3 repeated washing steps were conducted that were completed by a final 1 min PBS-T rinse. In a first step, slides were washed once with PBS-T to remove non-immobilized LPS and blocked with skim milk for 7 h. Following two washing steps, breast milk samples diluted to an IgA concentration of 10 μg/mL in blocking buffer were incubated for 16 h. The washing steps were repeated twice before the second antibody was added for 1 h incubation time. The secondary antibodies goat anti-human IgA Cy3 (109-165-011), donkey anti-human IgM Cy3 (709-165-073) and donkey anti-human IgG Alexa Fluor 647 (709-605-098) (all from Jackson ImmunoResearch, West Grove, PA, USA) were diluted 1:10,000 in blocking buffer. Washing steps were repeated twice followed by washing with 0.01% Tween 20 for 90 s. Finally, the slides were dried in the hybridization station HS4800 with nitrogen for 3.5 min at 30 °C. The slides were scanned immediately after processing using an LS400 confocal microarray scanner (Tecan, Männedorf, Switzerland). Cy3 fluorescence was excited at 532 nm and detected at 590 nm, Alexa Fluor 647 at 633nm. Mean net intensities in relative fluorescence units (RFU) and standard deviation (SD) were calculated using the ArrayPro 4.5 software (Media Cybernetics, Rockville, MD, USA).

### Leishmania lysates

*L. major* MHOM/SU/73/5ASKH and glf^−^ derivative (33) were grown at 27 °C in M199 medium (Invitrogen, Karlsruhe, Germany) supplemented with 10 % fetal calf serum, 40 mM Hepes pH 7.5, 0.1 mM adenine, 0.0005 % hemin, and 0.0002 % biotin. Exponentially growing promastigotes were harvested by centrifugation at 1300 x g for 10 min, washed with PBS, and resuspended at a concentration of 109 parasites/mL in 50 mM Tris/HCl pH8.0, 0.1% TritonX-100, 1mM phenylmethylsulfonyl fluoride (PMSF), 2 µg/mL Leupeptin and 5 µg/ml pepstatin. The parasites were homogenized by sonification, centrifuged at 1,300 x g for 10 min and the supernatant was collected. The protein concentration was determined in triplicate with a PierceTM BCA assay kit according to the manufacturer instructions. Laemmli buffer was added to the lysate and heated at 95 °C for 5 min.

### Plasmodium lysates

*P. falciparum* strain 3D7 parasites were cultured with human B+ erythrocytes (3 % hematocrit) in RPMI medium supplemented with Albumax and incubated at 37 °C in an atmosphere of 92 % N_2_, 3 % O_2_ and 5 % CO_2_ using standard methods (46). Human erythrocytes were purchased from the Banc de Sang i Teixits (Catalonia, Spain) after approval from the Comitè Ètic Investigació Clínica Hospital Clínic de Barcelona (HCB/2020/0051). Parasite growth was regularly monitored by counting the infected erythrocytes in Giemsa-stain blood smears by light microscopy. Asynchronous cultures at asexual intraerythrocytic stages with > 6-8 % parasitemia were centrifuged for 5 min at 1,500 rpm and the pellets resuspended in 2 red blood cell (RBC) volumes of 0.2 % saponin in PBS to disrupt RBCs membranes. The suspensions were incubated for 10 min on ice, then 10 mL of PBS were added and centrifuged for 8 min at 1,800 rpm at 4 °C. Supernatant was removed and saponin lysis was repeated. Uninfected B+ RBCs were used as controls. After centrifugation, the pellet was washed with PBS, transferred to a 1.5 mL tube and centrifuged at 10,000 x g for 10 min at 4 °C. Pellets were kept at −80 °C before total protein extraction. Frozen pellets were thoroughly resuspended with 200-300 μL Laemmli buffer (depending on the total number of parasites) and sonicated on ice three times for 10 s. Suspensions were then centrifuged at 10,000 x g for 30 min at 4 °C. Supernatants containing the extracted soluble proteins were stored at −80 °C before Western blot analysis.

### Immunofluorescence

*L. major* promastigotes were pelleted at 1,500 x g for 10 min washed and resuspended in PBS. Parasites were deposited on polylysine-coated coverslip and fixed with 4% paraformaldehyde for 15 min at room temperature. Pre-incubation, antibody incubation and washes were conducted in PBS buffer containing 2% bovine serum albumin (BSA). Polyclonal antisera specific to *E. coli* O55, *E. coli* O85, *E. coli* O142 and *E. coli* O152 O-antigens were purchased from SSI Diagnostica (Hillerød, Denmark). The antisera and Alexa Fluor488 goat anti-rabbit IgG1 (Jackson ImmunoResearch, West Grove, PA) were diluted 1:10 and 1:2,000, respectively. Nuclei were stained with 2.5 µg/ml 4′,6-diamidino-2-phenylindole. Coverslips were analyzed under a Zeiss Axiovert 200M microscope equipped with ApoTome module, AxioCam MRm digital camera, and Axio Vision software (Zeiss, Oberkochen, Germany). *P. falciparum* cultures were pelleted at 800 x g for 30 s, washed and resuspended in PBS. The culture was distributed on Concanavalin A-coated Ibidi removable wells and fixed with 4% paraformaldehyde for 10 min at room temperature. Autofluorescence was reduced by quenching with 0.1 M glycine in PBS for 10 min. RBCs were permeabilized with Triton X-100 0.2% PBS for 20 min and blocked with 5% BSA Triton X-100 0.2% PBS for 20 minutes at room temperature. Antibody incubation, diluted 1:10, and washes were conducted in 2% BSA and 0.2% Triton X-100 in PBS. The Alexa Fluor488 goat anti-rabbit IgG1 secondary antibody was diluted 1:500. A 2 µg/mL dilution of the polyclonal rabbit anti-PfHSP70 (StressMarq Bioscience, Canada) antibody was used as the positive control. Nuclei were stained with Hoechst 33342 and RBC surfaces with wheat germ agglutinin. Coverslips were analyzed under a Leica DMI4000 B microscope equipped with Leica CTR6500, Leica TCS SPE-II and LAS-AF Lite software (Leica, Wetzlar, Germany).

### Gel electrophoresis and Western blotting

LPS samples (15 µg) and parasite extracts (20 µg protein) were separated on 14 % acrylamide gels at 80-120 V for up to 3 h. Gels were either stained with silver (47)(48) and colloidal Coomassie (49) or used for Western blotting. For silver staining, gels were oxidized in 40 % ethanol, 5 % acetic acid, 0.7 % periodic acid in deionized water, followed by two washing steps of 20 min in 30 % ethanol and one washing step with deionized water for 20 min. Gels were sensitized for one minute in 0.02 % sodium thiosulfate in deionized water, washed three times for 20 s in deionized water before being stained with a 0.1 % silver nitrate solution for 20 min at 4 °*C.* Gels were washed three more times for 20 s in deionized water and finally developed in a solution containing 3 % sodium carbonate, 0.05 % formaldehyde in deionized water for around 5 min until bands appeared. After another washing step of 20 s with water, the reaction was stopped with 5 % acetic acid. Gels were stored in a 1 % acetic acid solution. For blotting, gels contents were transferred onto PDVF membranes at 250 mA for 1 h. Membranes were blocked overnight at 4 °C in 5 % skim milk solution in 1 x PBS-T supplemented with 0.1 % Tween 20. Polyclonal antisera specific to O-antigens from *E. coli* O55*, E. coli* O85, *E. coli* O142 and *E. coli* O152 were diluted 1:100 in PBS-T and incubated overnight at 4 °*C.* Membranes were washed 4-times for 10 min in 1 x PBS-T before 1 h incubation at room temperature with secondary anti-rabbit IgG coupled to horseradish peroxidase (Cell Signaling Technology, Danvers,MA, USA) diluted 1:1,000 in 1 x PBS-T. Membranes were washed for up to six times for 10 min with PBS-T, and antibody binding was detected using SuperSignal R West Pico PLUS chemiluminescence substrate (Thermo Fisher Scientific, Waltham, MA) and the FUSION FX imager (Vilber, Collégien, France).

### Statistical analysis

All statistical analysis were done with R studio version 4.1.1 (2021-08-10). Heatmaps with hierarchical clustering were done using pheatmaps (v1.0.12), heatmaps without clustering were done with ggplot2 (v3.4.1). The significance of IgA-LPS reactivity in Kenyan vs. Swiss samples was determined with the ggsignif package (v0.6.4) using the non-parametric and unpaired Wilcoxon test. Correlation between IgA/IgM or IgA/IgG binding towards certain LPS was tested using the Spearman correlation test. P-values ≤ 0.05 were considered statistically significant. The threshold of detection (TOD) at a RFU value of 88 was based on the signal intensity measured for empty samples and buffer controls.

## Author contributions

L.C. and T.H. conceived the study and designed the experiments. T.R. and D.B. obtained Swiss breast milk samples, D.P. and M.B.Z. obtained Kenyan breast milk samples. L.C. and N.M. performed ELISA for antibody determinations, isolated bacterial LPS and validated LPS integrity. Microarrays were printed by J.S, L.C. and N.M. performed microarray experiments. L.C. performed immunoblotting. P.Z., F.H.R, T.R.U and L.I. carried out parasite analysis. L.C. performed all data visualizations and statistical analysis. L.C. and T.H. wrote the initial draft of the manuscript. The manuscript was read by all authors and all comments contributed to the final draft.

## Acknowledgements

We thank Miriam Ramírez for producing the Plasmodium falciparum extracts.

## Data Availability

Datasets are available on request: The raw data supporting the conclusions of this article will be made available by the authors, without undue reservation.

## REFERENCES

1. Carr LE, Virmani MD, Rosa F, Munblit D, Matazel KS, Elolimy AA, et al. Role of Human Milk Bioactives on Infants’ Gut and Immune Health. Front Immunol. 2021;12:604080.

2. Hennet T, Borsig L. Breastfed at Tiffany’s. Trends in biochemical sciences. 2016;41(6):508–18.

3. Lyons KE, Ryan CA, Dempsey EM, Ross RP, Stanton C. Breast Milk, a Source of Beneficial Microbes and Associated Benefits for Infant Health. Nutrients. 2020;12(4).

4. Zheng W, Zhao W, Wu M, Song X, Caro F, Sun X, et al. Microbiota-targeted maternal antibodies protect neonates from enteric infection. Nature. 2020;577(7791):543–8.

5. Wilson E, Butcher EC. CCL28 controls immunoglobulin (Ig)A plasma cell accumulation in the lactating mammary gland and IgA antibody transfer to the neonate. J Exp Med. 2004;200(6):805–9.

6. Gopalakrishna KP, Macadangdang BR, Rogers MB, Tometich JT, Firek BA, Baker R, et al. Maternal IgA protects against the development of necrotizing enterocolitis in preterm infants. Nature Medicine. 2019;25(7):1110–5.

7. Durand D, Ochoa TJ, Bellomo SME, Contreras CA, Bustamante VH, Ruiz J, et al. Detection of Secretory Immunoglobulin A in Human Colostrum as Mucosal Immune Response Against Proteins of the Type III Secretion System of Salmonella, Shigella and Enteropathogenic Escherichia Coli. The Pediatric Infectious Disease Journal. 2013;32(10):1122–6.

8. Castro-Dopico T, Clatworthy MR. IgG and Fcγ Receptors in Intestinal Immunity and Inflammation. Front Immunol. 2019;10:805.

9. Rogier EW, Frantz AL, Bruno MEC, Wedlund L, Cohen DA, Stromberg AJ, et al. Secretory antibodies in breast milk promote long-term intestinal homeostasis by regulating the gut microbiota and host gene expression. Proceedings of the National Academy of Sciences. 2014;111(8):3074–9.

10. Mantis NJ, Rol N, Corthésy B. Secretory IgA’s complex roles in immunity and mucosal homeostasis in the gut. Mucosal Immunology. 2011;4(6):603–11.

11. Atyeo C, Alter G. The multifaceted roles of breast milk antibodies. Cell. 2021;184(6):1486–99.

12. Sterlin D, Fadlallah J, Adams O, Fieschi C, Parizot C, Dorgham K, et al. Human IgA binds a diverse array of commensal bacteria. J Exp Med. 2020;217(3).

13. Mostowy RJ, Holt KE. Diversity-Generating Machines: Genetics of Bacterial Sugar-Coating. Trends Microbiol. 2018;26(12):1008–21.

14. Cerutti A. The regulation of IgA class switching. Nature Reviews Immunology. 2008;8(6):421–34.

15. Pabst O, Slack E. IgA and the intestinal microbiota: the importance of being specific. Mucosal Immunology. 2020;13(1):12–21.

16. Kappler K, Hennet T. Emergence and significance of carbohydrate-specific antibodies. Genes and immunity. 2020;21(4):224–39.

17. Yilmaz B, Portugal S, Tran TM, Gozzelino R, Ramos S, Gomes J, et al. Gut microbiota elicits a protective immune response against malaria transmission. Cell. 2014;159(6):1277–89.

18. Latov N. Campylobacter jejuni Infection, Anti-Ganglioside Antibodies, and Neuropathy. Microorganisms. 2022;10(11):2139.

19. Raetz CRH, Whitfield C. Lipopolysaccharide endotoxins. Annual review of biochemistry. 2002;71:635–700.

20. Whitfield C, Williams DM, Kelly SD. Lipopolysaccharide O-antigens-bacterial glycans made to measure. J Biol Chem. 2020;295(31):10593–609.

21. Knirel YA, Valvano MA. Bacterial Lipopolysaccharides. Vienna: Springer Vienna; 2011. 444 p.

22. Giorgetti A, Paganini D, Nyilima S, Kottler R, Frick M, Karanja S, et al. The effects of 2’-fucosyllactose and lacto-N-neotetraose, galacto-oligosaccharides, and maternal human milk oligosaccharide profile on iron absorption in Kenyan infants. The American journal of clinical nutrition. 2023;117(1):64–72.

23. Jiang N, Dillon FM, Silva A, Gomez-Cano L, Grotewold E. Rhamnose in plants - from biosynthesis to diverse functions. Plant science : an international journal of experimental plant biology. 2021;302:110687.

24. Mistou M-Y, Sutcliffe IC, van Sorge NM. Bacterial glycobiology: rhamnose-containing cell wall polysaccharides in Gram-positive bacteria. FEMS Microbiology Reviews. 2016;40(4):464–79.

25. Adibekian A, Stallforth P, Hecht M-L, Werz DB, Gagneux P, Seeberger PH. Comparative bioinformatics analysis of the mammalian and bacterial glycomes. Chemical Science. 2011;2(2):337–44.

26. Smith SI, Seriki A, Ajayi A. Typhoidal and non-typhoidal Salmonella infections in Africa. Eur J Clin Microbiol Infect Dis. 2016;35(12):1913–22.

27. Nataro JP, Kaper JB. Diarrheagenic Escherichia coli. Clinical microbiology reviews. 1998;11(1):142–201.

28. Galili U, Rachmilewitz EA, Peleg A, Flechner I. A unique natural human IgG antibody with anti-alpha-galactosyl specificity. The Journal of Experimental Medicine. 1984;160:1519–31.

29. Oyelaran O, McShane LM, Dodd L, Gildersleeve JC. Profiling human serum antibodies with a carbohydrate antigen microarray. Journal of Proteome Research. 2009;8(9):4301–10.

30. Tefsen B, Ram AF, van Die I, Routier FH. Galactofuranose in eukaryotes: aspects of biosynthesis and functional impact. Glycobiology. 2011;22(4):456–69.

31. Njuguna P, Maitland K, Nyaguara A, Mwanga D, Mogeni P, Mturi N, et al. Observational study: 27 years of severe malaria surveillance in Kilifi, Kenya. BMC Medicine. 2019;17(1):124.

32. Ngere I, Gufu Boru W, Isack A, Muiruri J, Obonyo M, Matendechero S, et al. Burden and risk factors of cutaneous leishmaniasis in a peri-urban settlement in Kenya, 2016. PLoS One. 2020;15(1):e0227697.

33. Kleczka B, Lamerz A-C, van Zandbergen G, Wenzel A, Gerardy-Schahn R, Wiese M, et al. Targeted gene deletion of Leishmania major UDP-galactopyranose mutase leads to attenuated virulence. The Journal of biological chemistry. 2007;282(14):10498–505.

34. McGuire MK, Randall AZ, Seppo AE, Järvinen KM, Meehan CL, Gindola D, et al. Multipathogen Analysis of IgA and IgG Antigen Specificity for Selected Pathogens in Milk Produced by Women From Diverse Geographical Regions: The INSPIRE Study. Front Immunol. 2020;11:614372.

35. Takahashi T, Yoshida Y, Hatano S, Sugita-Konishi Y, Igimi S, Yajima M, et al. Reactivity of secretory IgA antibodies in breast milk from 107 Japanese mothers to 20 environmental antigens. Biology of the neonate. 2002;82(4):238–42.

36. Bunker JJ, Bendelac A. IgA Responses to Microbiota. Immunity. 2018;49(2):211–24.

37. Donald K, Petersen C, Turvey SE, Finlay BB, Azad MB. Secretory IgA: Linking microbes, maternal health, and infant health through human milk. Cell host & microbe. 2022;30(5):650–9.

38. Bunker JJ, Erickson SA, Flynn TM, Henry C, Koval JC, Meisel M, et al. Natural polyreactive IgA antibodies coat the intestinal microbiota. Science. 2017;358(6361).

39. Rollenske T, Szijarto V, Lukasiewicz J, Guachalla LM, Stojkovic K, Hartl K, et al. Cross-specificity of protective human antibodies against Klebsiella pneumoniae LPS O-antigen. Nature immunology. 2018;19(6):617–24.

40. Goyal PK, Wheatcroft J, Wakelin D. Tyvelose and protective responses to the intestinal stages of Trichinella spiralis. Parasitology international. 2002;51(1):91–8.

41. Turco SJ, Orlandi PA, Jr., Homans SW, Ferguson MA, Dwek RA, Rademacher TW. Structure of the phosphosaccharide-inositol core of the Leishmania donovani lipophosphoglycan. J Biol Chem. 1989;264(12):6711–5.

42. Haynes PA, Ferguson MA, Cross GA. Structural characterization of novel oligosaccharides of cell-surface glycoproteins of Trypanosoma cruzi. Glycobiology. 1996;6(8):869–78.

43. Cabezas-Cruz A, Mateos-Hernández L, Alberdi P, Villar M, Riveau G, Hermann E, et al. Effect of blood type on anti-α-Gal immunity and the incidence of infectious diseases. Exp Mol Med. 2017;49(3):e301.

44. Aguilar R, Ubillos I, Vidal M, Balanza N, Crespo N, Jiménez A, et al. Antibody responses to α-Gal in African children vary with age and site and are associated with malaria protection. Scientific reports. 2018;8(1):9999.

45. Davis MR, Jr., Goldberg JB. Purification and visualization of lipopolysaccharide from Gram-negative bacteria by hot aqueous-phenol extraction. J Vis Exp. 2012(63).

46. Trager W, Jensen JB. Human malaria parasites in continuous culture. Science (New York, NY). 1976;193(4254):673–5.

47. Blum H, Beier H, Gross HJ. Improved silver staining of plant proteins, RNA and DNA in polyacrylamide gels. Electrophoresis. 1987;8(2):93–9.

48. Fomsgaard A, Freudenberg MA, Galanos C. Modification of the silver staining technique to detect lipopolysaccharide in polyacrylamide gels. Journal of clinical microbiology. 1990;28(12):2627–31.

49. Dyballa N, Metzger S. Fast and sensitive colloidal coomassie G-250 staining for proteins in polyacrylamide gels. Journal of Visualized Experiments : JoVE. 2009(30).

50. Varki A, Cummings RD, Aebi M, Packer NH, Seeberger PH, Esko JD, et al. Symbol Nomenclature for Graphical Representations of Glycans. Glycobiology. 2015;25(12):1323–4.

