## Supplementary Material for "Inter-individual and inter-regional variability of breast milk antibody reactivity to bacterial lipopolysaccharides"

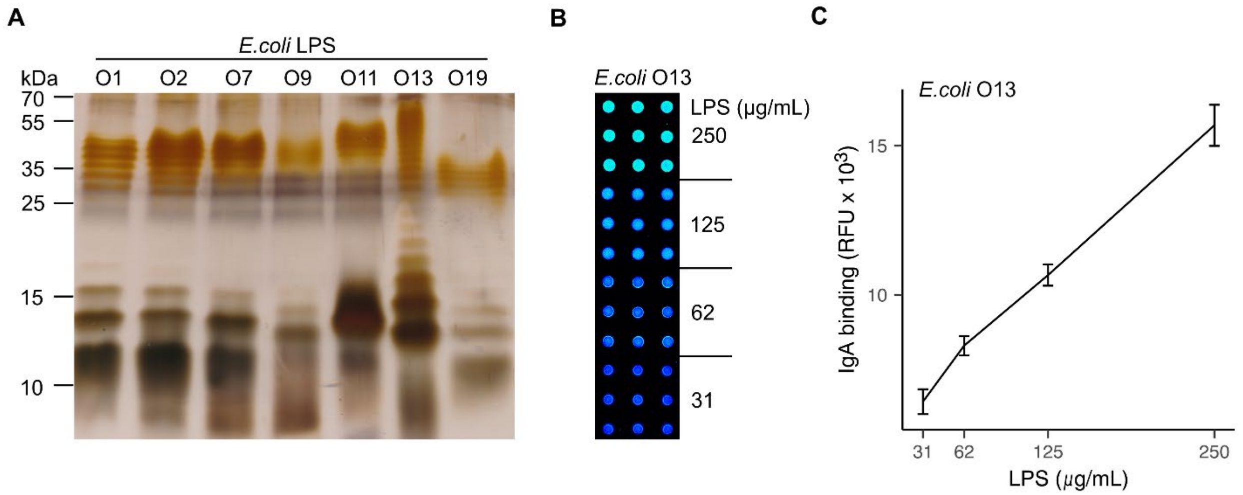


**Supplementary Figure 1.** LPS isolation and array printing. (A) SDS-PAGE and silver staining of LPS isolated from *E. coli serotypes*. (B) Fluorescence signal of *E. coli* O13 LPS printed in 9-dot matrices at 31, 62, 125 and 250 µg/ml visualized by scanning at 543 nm after binding to breast milk IgA and Cy3-labelled anti-human IgA antibody. (C) Quantitation of IgA binding to *E. coli* O13 LPS as mean ± S.D. of net fluorescence intensity from nine replicates per LPS concentration.

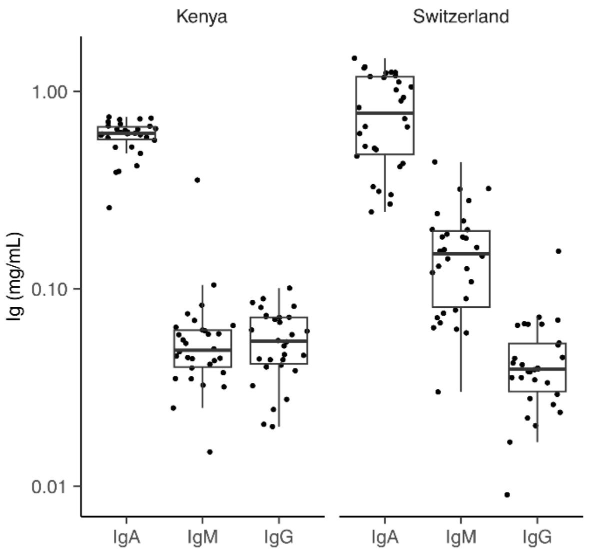


**Supplementary Figure 2.** Immunoglobulin concentrations in breast milk samples. IgA, IgM and IgG levels (mg/mL) in breast milk samples from Kenya and Switzerland were determined by ELISA. Unpaired Wilcoxon test used. IgM concentrations in samples from Switzerland were significantly higher than in Kenyan samples (p-value = 5.2x $10^{9-}$).

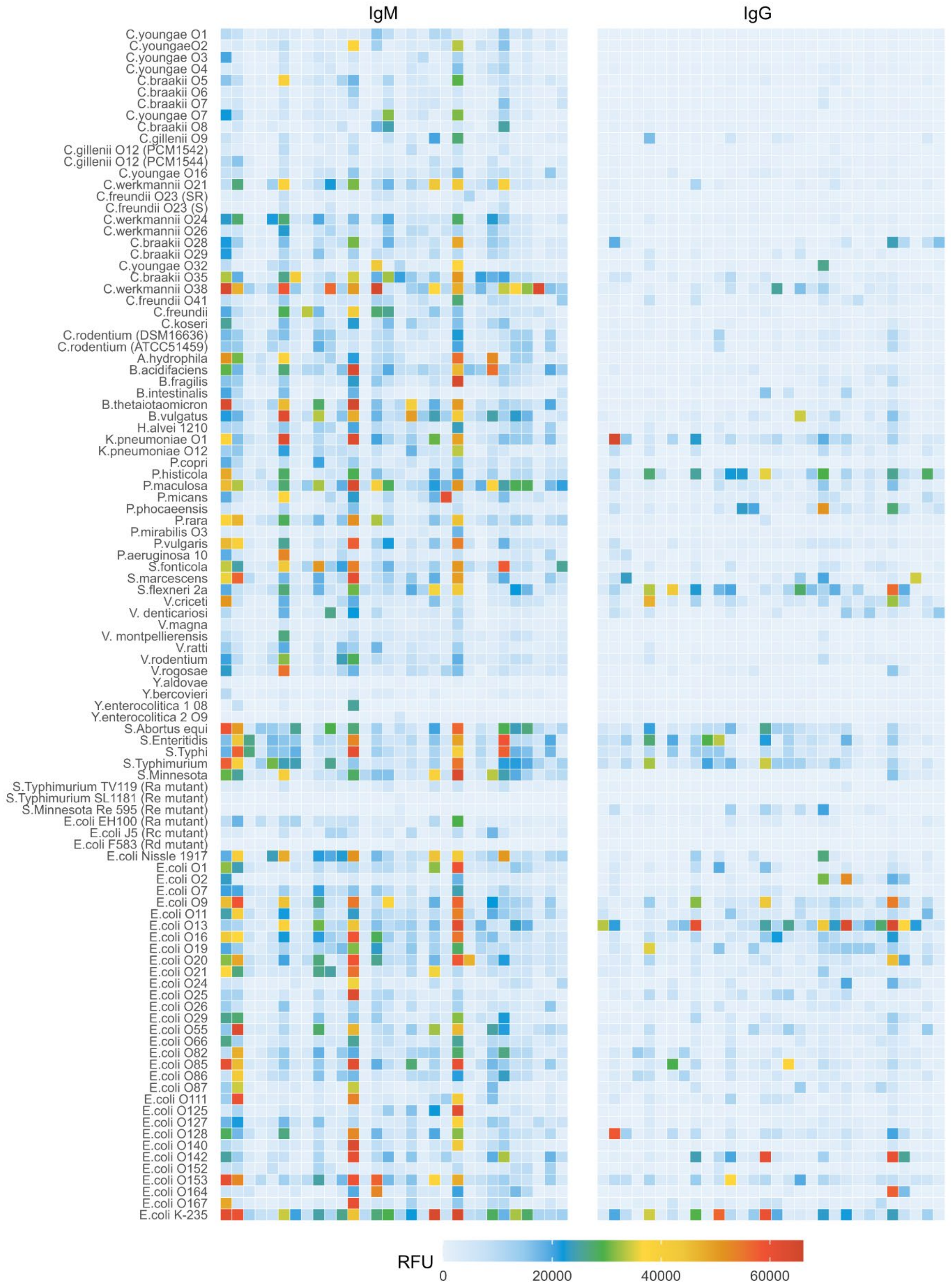
IgM
IgG
C.youngae O1
C.youngaeO2
C.youngae O3
C.youngae O4
C.braakii O5
C.braakii O6
C.braakii O7
C.youngae O7
C.braakii O8
C.gillenii O9
C.gillenii O12 (PCM1542)
C.gillenii O12 (PCM1544)
C.youngae O16
C.werkmannii O21
C.freundii O23 (SR)
C.freundii O23 (S)
C.werkmannii O24
C.werkmannii O26
C.braakii O28
C.braakii O29
C.youngae O32
C.braakii O35
C.werkmannii O38
C.freundii O41
C.freundii
C.koseri
C.rodentium (DSM16636)
C.rodentium (ATCC51459)
A.hydrophila
B.acidifaciens
B.fragilis
B.intestinalis
B.thetaiotaomicron
B.vulgatus
H.alvei 1210
K.pneumoniae O1
K.pneumoniae O12
P.copri
P.histicola
P.maculosa
P.micans
P.phocaeensis
P.rara
P.mirabilis O3
P.vulgaris
P.aeruginosa 10
S.fonticola
S.marcescens
S.flexneri 2a
V.criceti
V. denticariosi
V.magna
V. montpellierensis
V.ratti
V.rodentium
V.rogosae
Y.aldovae
Y.bercovieri
Y.enterocolitica 1 08
Y.enterocolitica 2 O9
S.Abortus equi
S.Enteritidis
S.Typhi
S.Typhimurium
S.Minnesota
S.Typhimurium TV119 (Ra mutant)
S.Typhimurium SL1181 (Re mutant)
S.Minnesota Re 595 (Re mutant)
E.coli EH100 (Ra mutant)
E.coli J5 (Rc mutant)
E.coli F583 (Rd mutant)
E.coli Nissle 1917
E.coli O1
E.coli O2
E.coli O7
E.coli O9
E.coli O11
E.coli O13
E.coli O16
E.coli O19
E.coli O20
E.coli O21
E.coli O24
E.coli O25
E.coli O26
E.coli O29
E.coli O55
E.coli O66
E.coli O82
E.coli O85
E.coli O86
E.coli O87
E.coli O111
E.coli O125
E.coli O127
E.coli O128
E.coli O140
E.coli O142
E.coli O152
E.coli O153
E.coli O164
E.coli O167
E.coli K-235
RFU
0
20000
40000
60000

**Supplementary Figure 3.** Breast milk IgM and IgG binding to LPS in Kenyan samples. Heatmap of IgM reactivity (left panel) detected using anti-human IgM Cy3 and scanned at 230 V PMT-gain. Heatmap of IgG reactivity (right panel) scanned at 250 V PMT-gain. Data are shown as net mean RFU values of 9-replicate dots.

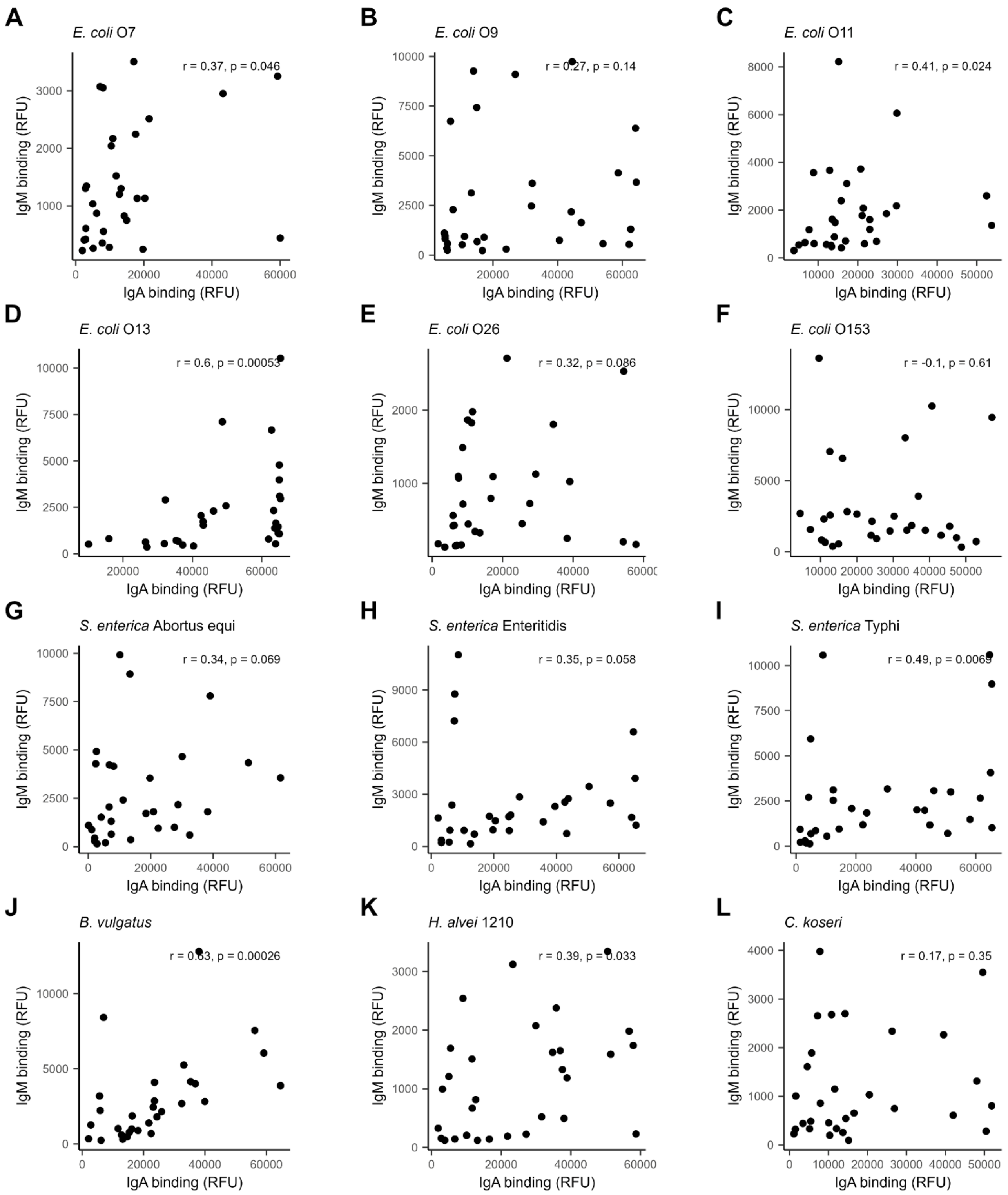


**Supplementary Figure 4.** Correlation of IgA and IgM binding to LPS. Data points represent individual breast milk samples from Kenya. IgA and IgM binding signals to specific LPS are given as net mean RFU. (A) *E. coli* O7, (B) *E. coli* O9, (C) *E. coli* O11, (D) *E. coli* O13, (E) *E. coli*

O26, (F)*E. coli* O153, (G) *S. enterica* Abortus equi, (H) *S. enterica* Enteriditis, (I) *S. enterica* Typhi, (J) *B. vulgatus,* (K) *H. alvei* 1210, (L) *C. koseri* LPS. Spearman correlation coefficients and p-values are listed for each panel.

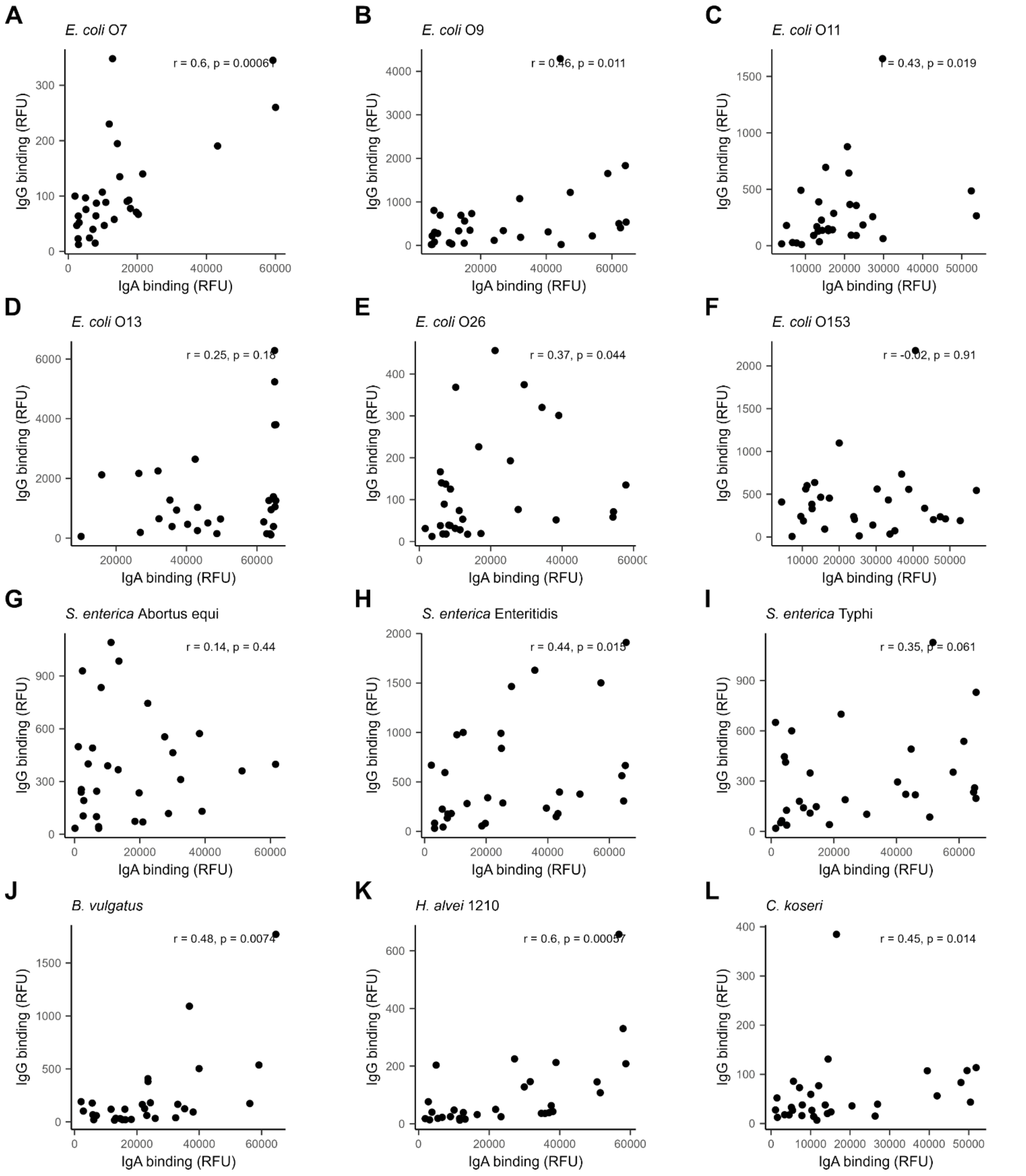


**Supplementary Figure 5.** Correlation of IgA and IgG binding to LPS. Data points represent individual breast milk samples from Kenya. IgA and IgG binding signals to specific LPS are given as net mean RFU. (A) *E. coli* O7, (B) *E. coli* O9, (C) *E. coli* O11, (D) *E. coli* O13, (E) *E. coli* O26, (F)

*E. coli* O153, (G) *S. enterica* Abortus equi, (H) *S. enterica* Enteriditis, (I) *S. enterica* Typhi, (J) *B. vulgatus,* (K) *H. alvei* 1210, (L) *C. koseri* LPS. Spearman correlation coefficients and p-values are listed for each panel.

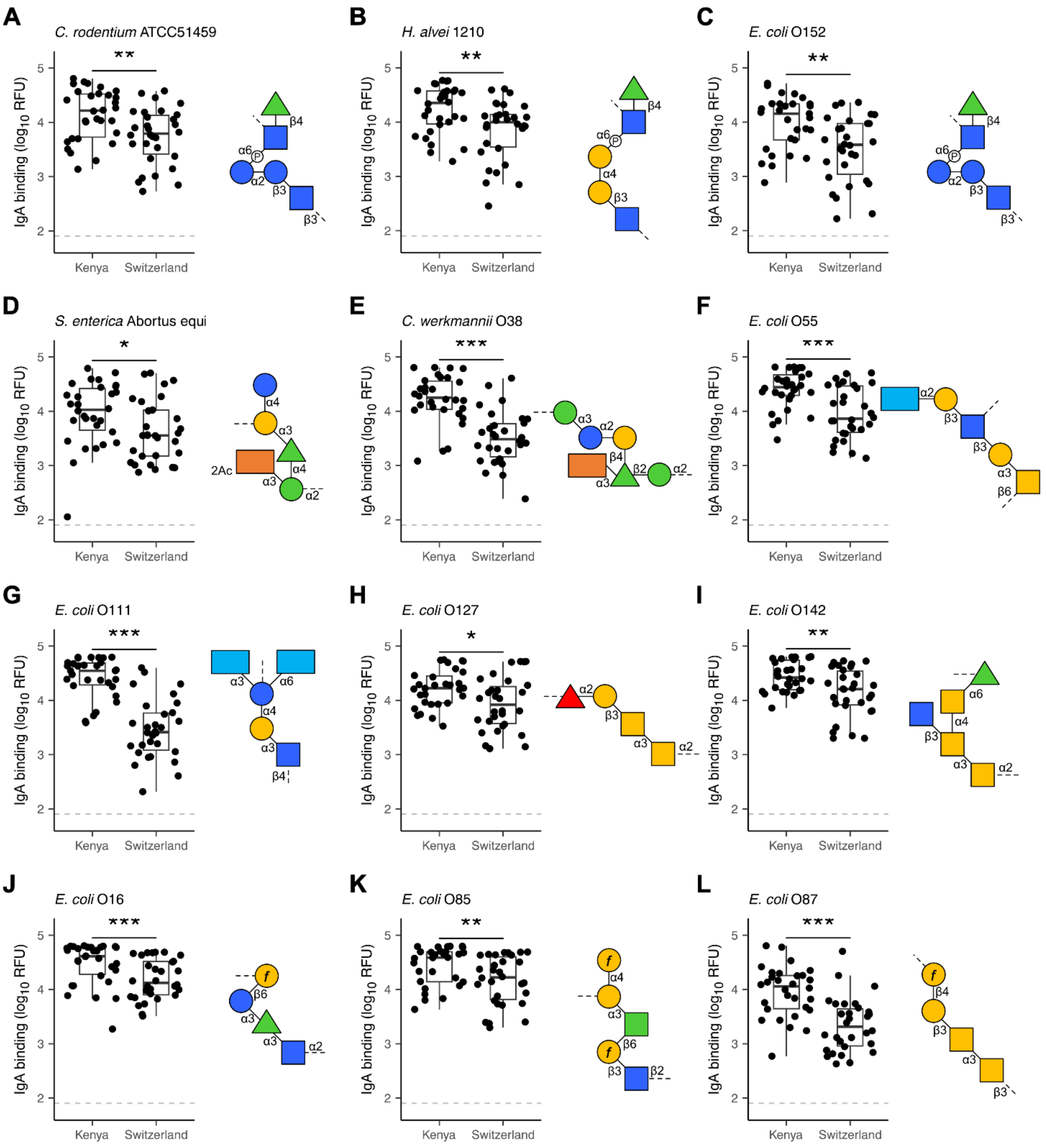


**Supplementary Figure 6.** Comparison of breast milk IgA reactivity towards specific LPS. LPS containing rhamnose (green triangle) as in (A) *C. rodentium*, (B) *H. alvei* 1210 and (C) *E. coli* O152 .LPS containing the 1,3-dideoxy sugars abequose (orange rectangle) and colitose (turquoise rectangle) as found in (D) *S. enterica* Abortus equi, (E) *C. werkmannii* O38, (F) *E. coli*

O55 and (G) *E. coli* O111 .LPS featuring the Forssman antigen (two yellow squares) as found in (H) *E. coli* O127, (I) *E. coli* O142, (L) *E. coli* O87. LPS containing galactofuranose (*f* in yellow circle) as in (J) *E. coli* O16, (K) *E. coli* O85 and (L) *E. coli* O87. IgA binding was detected using anti-human IgA Cy3, scanned at PMT-gain 190V. Comparisons of net mean RFU values between Kenyan (n=30) and Swiss (n=30) samples using unpaired Wilcoxon test. P-values ≤ 0.05 were considered significant ("***"=0.001, "**"=0.01, "*"=0.05). The grey dotted line marks the threshold of detection (TOD).

**Supplementary Table 1.** LPS library printed on arrays. Pure bacterial cultures were grown under recommended conditions (medium, temperature and oxygen requirements) by the supplier. No growth media are indicated for commercially supplied LPS. Known O-antigen structures are referenced.

| O-antigen | Source | Culture conditions | | | Structure |
|---|---|---|---|---|---|
| | | Growth medium | Temperature | Oxygen | |
| ***Aeromonas hydrophila*** | DSM 30187 | Trypticase Soy Yeast Extract Medium (DSMZ 92) | 30°C | aerobic | N/A |
| ***Bacteroides acidifaciens*** | DSM 100502 | Chopped Meat Medium (DSMZ 78) | 37°C | anaerobic | N/A |
| ***Bacteroides fragilis*** | DSM 2151 | Chopped Meat Medium (DSMZ 78) | 37°C | anaerobic | N/A |
| ***Bacteroides intestinalis*** | DSM 17393 | modified PYG medium (DSMZ 104) | 37°C | anaerobe | N/A |
| ***Bacteroides thetaiotaomicron*** | DSM 2079 | Chopped Meat Medium (DSMZ 78) + haemin and vitamin K1 | 37°C | anaerobe | N/A |
| ***Bacteroides vulgatus*** | DSM 1447 | Chopped Meat Medium (DSMZ 78) + haemin and vitamin K1 | 37°C | anaerobe | N/A |
| ***Citrobacter youngae* O1** | PCM 1506 | Nutrient broth (DSMZ 1) | 37°C | aerobe | [1] |

| | | | | | |
|---|---|---|---|---|---|
| ***Citrobacter youngae* O2** | PCM 1507 | Nutrient broth (DSMZ 1) | 37°C | aerobe | [1] |
| ***Citrobacter youngae* O3** | PCM 1509 | Nutrient broth (DSMZ 1) | 37°C | aerobe | [1] |
| ***Citrobacter youngae* O4** | PCM 1525 | Nutrient broth (DSMZ 1) | 37°C | aerobe | [1] |
| ***Citrobacter braakii* O5** | PCM 1528 | Nutrient broth (DSMZ 1) | 37°C | aerobe | [1] |
| ***Citrobacter braakii* O6** | PCM 1531 | Nutrient broth (DSMZ 1) | 37°C | aerobe | [1] |
| ***Citrobacter braakii* O7** | PCM 1532 | Nutrient broth (DSMZ 1) | 37°C | aerobe | [1] |
| ***Citrobacter youngae* O7** | PCM 1503 | Nutrient broth (DSMZ 1) | 37°C | aerobe | [1] |
| ***Citrobacter braakii* O8** | PCM 1536 | Nutrient broth (DSMZ 1) | 37°C | aerobe | [1] |
| ***Citrobacter gillenii* O9** | PCM 1537 | Nutrient broth (DSMZ 1) | 37°C | aerobe | [1] |
| ***Citrobacter gillenii* O12** | PCM 1544 | Nutrient broth (DSMZ 1) | 37°C | aerobe | [1] |
| ***Citrobacter gillenii* O12** | PCM 1542 | Nutrient broth (DSMZ 1) | 37°C | aerobe | [1] |

| | | | | | |
|---|---|---|---|---|---|
| ***Citrobacter youngae* O16** | PCM 1550 | Nutrient broth (DSMZ 1) | 37°C | aerobe | [1] |
| ***Citrobacter werkmanii* O21** | PCM 1554 | Nutrient broth (DSMZ 1) | 37°C | aerobe | [1] |
| ***Citrobacter freundii* O23 (SR)** | PCM 1556 | Nutrient broth (DSMZ 1) | 37°C | aerobe | [1] |
| ***Citrobacter freundii* O23 (S)** | PCM 2352 | Nutrient broth (DSMZ 1) | 37°C | aerobe | [1] |
| ***Citrobacter werkmanii* O24** | PCM 1557 | Nutrient broth (DSMZ 1) | 37°C | aerobe | [1] |
| ***Citrobacter werkmanii* O26** | PCM 1559 | Nutrient broth (DSMZ 1) | 37°C | aerobe | [1] |
| ***Citrobacter braakii* O28** | PCM 1561 | Nutrient broth (DSMZ 1) | 37°C | aerobe | [1] |
| ***Citrobacter braakii* O29** | PCM 1562 | Nutrient broth (DSMZ 1) | 37°C | aerobe | [1] |
| ***Citrobacter youngae* O32** | PCM 1569 | Nutrient broth (DSMZ 1) | 37°C | aerobe | [1] |
| ***Citrobacter braakii* O35** | PCM 1586 | Nutrient broth (DSMZ 1) | 37°C | aerobe | [1] |
| ***Citrobacter werkmanii* O38** | PCM 1489 | Nutrient broth (DSMZ 1) | 37°C | aerobe | [1] |

| | | | | | |
|---|---|---|---|---|---|
| ***Citrobacter freundii* O41** | PCM 1444 | Nutrient broth (DSMZ 1) | 37°C | aerobe | [1] |
| ***Citrobacter freundii*** | DSM 30039 | Trypticase Soy Yeast Extract Medium (DSMZ 92) | 30°C | aerobe | N/A |
| ***Citrobacter koseri*** | DSM 4595 | Nutrient broth (DSMZ 1) | 30°C | aeroba | N/A |
| ***Citrobacter rodentium*** | DSM 16636 | Nutrient broth (DSMZ 1) | 37°C | aeroba | N/A |
| ***Citrobacter rodentium*** | ATCC 51459 | LB broth (DSMZ 381) | 37°C | aerobe | [1] |
| ***Escherichia coli* EH100** | L9641 | - | - | - | Ra rough mutant |
| ***Escherichia coli* J5** | L5014 | - | - | - | Rc rough mutant |
| ***Escherichia coli* F583** | L6893 | - | - | - | Rd rough mutant |
| ***Escherichia coli* K-235** | L2143 | - | - | - | N/A |
| ***Escherichia coli* Nissle 1917** | | LB broth (DSMZ 381) | 37°C | aerobe | [2] |
| ***Escherichia coli* O1** | DSM 10728 | LB broth (DSMZ 381) | 37°C | aerobe | [3] |

| | | | | | |
|---|---|---|---|---|---|
| ***Escherichia coli* O2** | DSM 10772 | LB broth (DSMZ 381) | 37°C | aerobe | [3] |
| ***Escherichia coli* O7** | DSM 10782 | LB broth (DSMZ 381) | 37°C | aerobe | [3] |
| ***Escherichia coli* O9** | NCTC 9009 | LB broth (DSMZ 381) | 37°C | aerobe | [3] |
| ***Escherichia coli* O11** | DSM 11751 | LB broth (DSMZ 381) | 37°C | aerobe | [3] |
| ***Escherichia coli* O13** | DSM 1328 | LB broth (DSMZ 381) | 37°C | aerobe | [3] |
| ***Escherichia coli* 016** | NCTC9016 | LB broth (DSMZ 381) | 37°C | aerobe | [3] |
| ***Escherichia coli* O19** | DSM 11752 | LB broth (DSMZ 381) | 37°C | aerobe | [3] |
| ***Escherichia coli* O20** | NCTC 9020 | LB broth (DSMZ 381) | 37°C | aerobe | [3] |
| ***Escherichia coli* O21** | NCTC9021 | LB broth (DSMZ 381) | 37°C | aerobe | [3] |
| ***Escherichia coli* O24** | NCTC 9024 | LB broth (DSMZ 381) | 37°C | aerobe | [3] |
| ***Escherichia coli* O25** | DSM 22664 | LB broth (DSMZ 381) | 37°C | aerobe | [3] |

| | | | | | |
|---|---|---|---|---|---|
| ***Escherichia coli* O26** | L8274 | - | - | - | [3] |
| ***Escherichia coli* O29** | DSM 9026 | LB broth (DSMZ 381) | 37°C | aerobe | [3] |
| ***Escherichia coli* O55** | L2880 | - | - | - | [3] |
| ***Escherichia coli* O66** | NCTC 9066 | LB broth (DSMZ 381) | 37°C | aerobe | [3] |
| ***Escherichia coli* O82** | NCTC 9082 | LB broth (DSMZ 381) | 37°C | aerobe | [3] |
| ***Escherichia coli* O85** | NCTC 9085 | LB broth (DSMZ 381) | 37°C | aerobe | [3] |
| ***Escherichia coli* O86** | NCTC 9086 | LB broth (DSMZ 381) | 37°C | aerobe | [3] |
| ***Escherichia coli* O87** | NCTC 9087 | LB broth (DSMZ 381) | 37°C | aerobe | [3] |
| ***Escherichia coli* O111** | L2630 | - | - | - | [3] |
| ***Escherichia coli* O125** | DSM 8700 | LB broth (DSMZ 381) | 37°C | aerobe | [3] |
| ***Escherichia coli* O127** | L3129 | - | - | - | [3] |

| | | | | | |
|---|---|---|---|---|---|
| ***Escherichia coli* O128** | L2755 | - | - | - | [3] |
| ***Escherichia coli* O140** | NCTC 10087 | LB broth (DSMZ 381) | 37°C | aerobe | [3] |
| ***Escherichia coli* O142** | DSM 9024 | LB broth (DSMZ 381) | 37°C | aerobe | [3] |
| ***Escherichia coli* O152** | DSM 9030 | LB broth (DSMZ 381) | 37°C | aerobe | [3] |
| ***Escherichia coli* O153** | NCTC 10960 | LB broth (DSMZ 381) | 37°C | aerobe | [3] |
| ***Escherichia coli* O164** | NCTC 10361 | LB broth (DSMZ 381) | 37°C | aerobe | [3] |
| ***Escherichia coli* O167** | DSM 9033 | LB broth (DSMZ 381) | 37°C | aerobe | [3] |
| ***Hafnia alvei* 1210** | PCM 1210 | Nutrient broth (DSMZ 1) | 37°C | aerobe | [4] |
| ***Klebsiella pneumoniae* O1** | L4268 | - | - | - | [5] |
| ***Klebsiella pneumoniae* O12** | PCM 80 | Nutrient broth (DSMZ 1) | 37°C | aerobe | [5] |
| ***Prevotella copri*** | DSM 18205 | Schaedler broth (Roth;5772) | 37°C | anaerobe | N/A |
| ***Prevotella histicola*** | DSM 19854 | modified PYG medium (DSMZ 104) | 37°C | anaerobe | N/A |

| | | | | | |
|---|---|---|---|---|---|
| ***Prevotella maculosa*** | DSM 19339 | modified PYG medium (DSMZ 104) | 37°C | anaerobe | N/A |
| ***Prevotella micans*** | DSM 21469 | modified PYG medium (DSMZ 104) + 5 % horse serum | 37°C | anaerobe | N/A |
| ***Prevotella phocaeensis*** | DSM 103364 | Chopped meat medium (DSMZ 78) + haemin and vitamin K1 | 37°C | anaerobe | N/A |
| ***Prevotella rara*** | DSM 105141 | Chopped meat medium (DSMZ 78) + haemin and vitamin K1 | 37°C | anaerobe | N/A |
| ***Prevotella salivae*** | DSM 15606 | Chopped meat medium (DSMZ 78) | 37°C | anaerobe | N/A |
| ***Proteus vulgaris*** | SMB00801 | - | - | - | N/A |
| ***Pseudomonas aeruginosa* 10** | L9143 | - | - | - | N/A |
| ***Salmonella enterica* Abortus equi** | L5886 | - | - | - | [6] |
| ***Salmonella enterica* Enteritidis** | L6011 | - | - | - | [6] |

| | | | | | |
|---|---|---|---|---|---|
| ***Salmonella enterica* Minnesota** | L6261 | - | - | - | N/A |
| ***Salmonella entérica* Minnesota Re 595** | L9764 | - | - | - | Re rough mutant |
| ***Salmonella enterica* Typhimurium** | L6511 | - | - | - | [6] |
| ***Salmonella enterica* Typhimurium SL1181** | L9516 | - | - | - | Re rough mutant |
| ***Salmonella enterica* Typhimurium TV119** | L6016 | - | - | - | Ra rough mutant |
| ***Salmonella enterica* Typhi** | L6386 | - | - | - | [6] |
| ***Serratia fonticola*** | DSM 103300 | Nutrient broth (DSMZ 1) | 28°C | aerobe | N/A |
| ***Serratia marcescens*** | L6136 | - | - | - | N/A |
| ***Shigella flexneri* 2a** | DSM 4782 | Trypticase Soy Yeast Extract Medium (DSMZ 92) | 37°C | aerobe | [7] |

| | | | | | |
|---|---|---|---|---|---|
| ***Veillonella criceti*** | DSM 20734 | Veillonella medium (DSMZ 136) | 37°C | anaerobe | N/A |
| ***Veillonella denticariosi*** | DSM 19009 | Veillonella medium (DSMZ 136) | 37°C | anaerobe | N/A |
| ***Veillonella magna*** | DSM 19857 | modified PYG medium (DSMZ 104) + 0.5 - 1% lactate | 37°C | anaerobe | N/A |
| ***Veillonella montpellierensis*** | DSM 17217 | modified PYG medium (DSMZ 104) + 10 µl/mL putrescine (0.03 %) + 10 µl/mL sodium lactate (50 %) | 37°C | anaerobe | N/A |
| ***Veillonella ratti*** | DSM 20736 | Veillonella medium (DSMZ 136) | 37°C | anaerobe | N/A |
| ***Veillonella rodentium*** | DSM 20737 | Veillonella medium (DSMZ 136) | 37°C | anaerobe | N/A |
| ***Veillonella rogosae*** | DSM 18960 | modified PYG medium (DSMZ 104) + 0.5 – 1 % lactate | 37°C | anaerobe | N/A |
| ***Yersinia aldovae*** | DSM 18303 | Trypticase Soy broth (Oxoid CM0129) | 30°C | aerobe | N/A |
| ***Yersinia bercovieri*** | DSM 18528 | Trypticase Soy broth (Oxoid CM0129) | 30°C | aerobe | N/A |

| | | | | | |
|---|---|---|---|---|---|
| ***Yersinia enterocolitica* O8** | DSM 27689 | BHI medium (Sigma-Aldrich; 53286) | 28°C | aerobe | [8] |
| ***Yersinia enterocolitica* O9** | DSM 9499 | BHI medium (Sigma-Aldrich; 53286) | 28°C | aerobe | [8] |


1. Knirel, Y.A., et al., *Structures and serology of the O-specific polysaccharides of bacteria of the genus Citrobacter.* Archivum immunologiae et therapiae experimentalis, 2002. **50**(6): p. 379–391.
2. Blum, G., R. Marre, and J. Hacker, *Properties of Escherichia coli strains of serotype O6.* Infection, 1995. **23**(4): p. 234–236.
3. Stenutz, R., A. Weintraub, and G. Widmalm, *The structures of Escherichia coli O-polysaccharide antigens.* FEMS microbiology reviews, 2006. **30**(3): p. 382–403.
4. Romanowska, E., *Immunochemical aspects of Hafnia alvei O antigens.* FEMS immunology and medical microbiology, 2000. **27**(3): p. 219–225.
5. Vinogradov, E., et al., *Structures of lipopolysaccharides from Klebsiella pneumoniae. Eluicidation of the structure of the linkage region between core and polysaccharide O chain and identification of the residues at the non-reducing termini of the O chains.* Journal of Biological Chemistry, 2002. **277**(28): p. 25070–25081.
6. Liu, B., et al., *Structural diversity in Salmonella O antigens and its genetic basis.* FEMS microbiology reviews, 2014. **38**(1): p. 56–89.
7. Perepelov, A.V., et al., *Shigella flexneri O-antigens revisited: final elucidation of the O-acetylation profiles and a survey of the O-antigen structure diversity.* FEMS Immunology & Medical Microbiology, 2012. **66**(2): p. 201–210.
8. Bruneteau, M. and S. Minka, *Lipopolysaccharides of bacterial pathogens from the genus Yersinia: a mini-review.* Biochimie, 2003. **85**(1-2): p. 145–152.

**Supplementary Table 2.** LPS Overview of breast milk samples. Breast milk samples used in this stud originated from Switzerland or Kenya, maternal age range and the IgA, IgM and IgG concentrations (mg/mL) determined with ELISA.

| Origin | Maternal age (years) | Antibody concentrations (mg/mL) | | |
|---|---|---|---|---|
| | | IgA | IgM | IgG |
| Kenya | 18-40 | 0.25 - 0.74 | 0.01 - 0.35 | 0.02 - 0.10 |
| Switzerland | 23-41 | 0.24 - 1.47 | 0.03 - 0.44 | 0.01 - 0.15 |
